# Multi-material volumetric bioprinting and plug-and-play suspension bath biofabrication via bioresin molecular weight tuning and multiwavelength optics

**DOI:** 10.1101/2024.09.21.614231

**Authors:** Davide Ribezzi, Jan-Philip Zegwaart, Thomas Van Gansbeke, Aitor Tejo-Otero, Sammy Florczak, Joska Aerts, Paul Delrot, Andreas Hierholzer, Martin Fussenegger, Jos Malda, Jos Olijve, Riccardo Levato

## Abstract

Volumetric Bioprinting (VBP), enables to rapidly build complex, cell-laden hydrogel constructs for tissue engineering and regenerative medicine. Light-based tomographic manufacturing enables spatial-selective polymerization of a bioresin, resulting in higher throughput and resolution than what achieved using traditional techniques. However, methods for multi-material printing are needed for a broad VBP adoption and applicability. Although converging VBP with extrusion bioprinting in support baths offers a novel, promising solution, further knowledge on the engineering of hydrogels as light-responsive, volumetrically printable baths is needed. Therefore, this study investigates the tuning of gelatin macromers, in particular leveraging the effect of molecular weight and degree of modification, to overcome these challenges, creating a library of materials for VBP and Embedded extrusion Volumetric Printing (EmVP). Bioresins with tunable printability and mechanical properties are produced, and a novel subset of gelatins and GelMA exhibiting stable shear-yielding behavior offers a new, single-component, ready-to-use suspension medium for in-bath printing, which is stable over multiple hours without needing temperature control. As proof-of-concept biological application, bioprinted gels are tested with insulin-producing pancreatic cell lines for 21 days of culture. Leveraging a multi-color printer, complex multi-material and multi-cellular geometries are produced, enhancing the accessibility of volumetric printing for advanced tissue models.

## 1. INTRODUCTION

Recent advances in 3D printing technologies led to the emergence of volumetric 3D printing, also termed volumetric additive manufacturing, a new family of techniques capable to produce objects of virtually any shape and size in a matter of seconds.^[1–3]^ With specific focus on the field of biofabrication, the development of Volumetric Bioprinting (VBP) enabled to sculpt hydrogels and living cells into centimeter-scale, clinically-relevant sized constructs at unprecedented velocity.^[3]^ This opens up to new avenues for large-scale tissue engineering and regenerative medicine, as well as for the high-throughput production of advanced *in vitro* tissue models for biomedical and pharmaceutical research.^[3,4]^ While several volumetric printing modalities have been introduced,^[5–8]^ the fastest approach to date to build convoluted porous geometries typical of living tissues, relies on the principles of tomographic manufacturing. In this respect, VBP is performed projecting visible light from a spatial light modulator (i.e. a digital micromirror device, DMD) onto a rotating vial containing a photoresponsive, cell-laden hydrogel, also termed bioresin. At each angle of rotation, the DMD delivers a different 2D projection, calculated following a tomographic reconstruction algorithm. Altogether, the light projections induce a cumulative light dose exceeding the threshold of crosslinking of the bioresin only in the voxels that correspond to the object to be printed, which solidifies in a layer-less fashion. This mode of printing results in much faster (<15 s) fabrication rates compared to layer-by-layer extrusion and digital light processing (DLP) or stereolithographic (SLA) printing, while achieving resolutions in the range of 40 µm, and allowing to safely process fragile cells and organoids in absence of mechanical stresses.^[4]^

Nevertheless, for VBP to be broadly adopted, key challenges remain to be overcome. First, tomographic printing introduces new requirements in terms of materials properties (optical, rheological) for the bioresins. In particular, accurate tuning of the photocrosslinking kinetics is paramount to produce accurate prints, avoid excessive light dose accumulation in non-desired regions of the bioresin vial. Moreover, methods for facile multi-material printing and for the production of complex tissues comprising multiple cell populations are needed. We recently introduced the concept of Embedded extrusion Volumetric Printing (EmVP), a technique converging suspended bath bioprinting, as a mean to position multiple cell types in a bioresin vat, and VBP to sculpt the now isotropic bioresin into any desired geometry. This method relied on the use of photoannelable microparticles (or microgels) as bioresin. This granular composirion, prior to photocrosslinking, act as a Bingham-like material suitable as bath for suspended printing.^[9]^ While versatile, this approach limits the printing resolution to the size of the microgels. Other groups later explored the use of gelatin as sacrificial, and thermoreversible viscosity enhancer blended with other resins of interest.^[10]^ However, this approach is challenged by limited printing resolution and a short time window of printability, since gelatin typically display and ideal temperature-dependent, shear thinning behaviour, restricted in a narrow (few minutes only) range during its thermal gelation curve^[10]^.

Herein, we propose that engineering the gelatin macromers will allow to overcome these challenges, and establish a library of fit-for-purpose materials for VBP, suspended bath printing and EmVP. Notably, gelatin, and particularly its photocrosslinkable derivative gelatin methacryloyl (GelMA), have become golden standard materials in the field of bioprinting. This is due to their availability in large amounts, biocompatibility, content of integrin-binding domains (i.e. RGD sequences), and intrinsic capacity to sustain cell adhesion and proliferation.^[11,12]^ Despite the relevant body of literature concerning these materials, key polymer phyisco-chemical properties, such as molecular weight,^[13–15]^ are mostly overlooked. If properly tuned, it could give rise to bioresins and hydrogels with broadly different characteristics, printability and biological performances. In this study, we investigate the design of bioresins obtained from gelatins with varying molecular weight (MW) and Degree of Modification (DoM) to produce a library of materials compatible with a multi-wavelength VBP printer and to identify groups of bioresins able to act as suspension baths printing and for EmVP. The effect of the synergy between MW, DoM, and temperature on the mechanical and rheological properties, and on the printability via VBP is studied. Bioresins suitable as mechanical support and as soft viscoelastic matrices for the culture of pancreatic cells are identified. Notably, a subset of gelatins and GelMA that exhibit shear-yielding behaviour in a stable manner over multiple hours was discovered, overcoming the limits of currently utilized gelatins, and offering a new material as ready-to-use suspension media for in-bath printing, with improved resolution over microgel-based methods. Finally, leveraging a multi-color optics VBP printer, well-aligned multi-material and multi-cellular, complex geometries were produced and demonstrated, by means of sequential VBP and EmVP. We expect that the ease of applicability of this method will extend the accessibility of volumetric printing for producing advanced, heterocellular tissue models for applications in tissue engineering and regenerative medicine.

## 2. RESULTS AND DISCUSSION

### 2.1 An “old” polymer under a new light: generating a library of gelatin-based hydrogels for VBP with tunable printability and mechanics

In VBP, printing accuracy relies primarily on the ability to precisely deliver spatially controlled light doses with the printing volume, and on the kinetics of crosslinking of the bioresin. The latter is naturally dependent not only on the light dose and photoinitiator concentration, but also on the available photoreactive groups (methacrylates, in the case of GelMA), which is a function of the molecular weight and degree of modification. Therefore, starting from type A porcine GelMA formulations that have average molecular weights of 90 and 160 kDa, with various degrees of modification (40, 60 or 80%), we generated a library of mechanical properties for the different GelMA hydrogels formulations. Error! Reference source not found.A depicts the photo-rheological data of GelMAs with a molar mass of 90 kDa and 160 kDa at DoM of 40, 60 and 80%, and at concentration of 5, 10, 15 and 20% w/v, in presence of lithium phenyl-2,4,6-trimethylbenzoylphosphinate (LAP) as photoinitiator. Regardless of the MW, all formulations rapidly crosslinked and reached a plateau or storage modulus within 8 minutes of illumination. The formulations that were expected to produce the densest hydrogel networks (concentration >15%, 80% DoM) displayed syneresis, the partial shrinking of the hydrogel induced by expelling water not bound to the tight polymeric mesh. This effect is visualized in the rheometer by an artefactual drop in storage modulus, caused by a loss of contact between the gel and the rheometer geometry upon shrinking.^[16,17]^ The crosslinking kinetics of GelMA-based hydrogels plays a pivotal role during light-based fabrication processes, as it affects the final mechanical properties and the printing resolution of the final model. Characterising the light dose needed to reach an optimal crosslinking condition is especially crucial when such hydrogels are employed for volumetric bioprinting. With this technique, since the whole bioresin volume is continuously photoexposed, excessive printing time can lead to off-target polymerization and loss of printing fidelity.^[1–3]^ For this reason, we characterised how the interplay between the molar mass and degree of modification translated to the optimal printing dose within our volumetric printing set up, a schematic of which is represented in Figure 1B, together with a proof-of-concept 3D print of a pancreas model. Results reported in Figure 1C showed how varying the MW and DoM lead to a different optimal dose for each studied formulation, resulting in 154.5 mJ/cm^2^, 117 mJ/cm^2^ and 142.5 mJ/cm^2^ for GelMA 90p80, 160p80 and 160p40, respectively (the three different GelMA formulations were chosen to have a side-by-side comparison of two formulations with different MW and DoM). These values correspond to a printing time of 16 s, 12 s, and 14 s. It should be noted that, during the VBP printing process, partial crosslinking of the gel is needed and reached, while complete crosslinking and consumption of the photoreactive groups can be obtained by post-curing after printing. Exceeding the optimal printing dose results in over-polymerized structures, as shown when all bioresin formulations were printed at the optimal, reference dose, identified for the 90p80 material (154.5 mJ/cm^2^).

**FIGURE 1:**
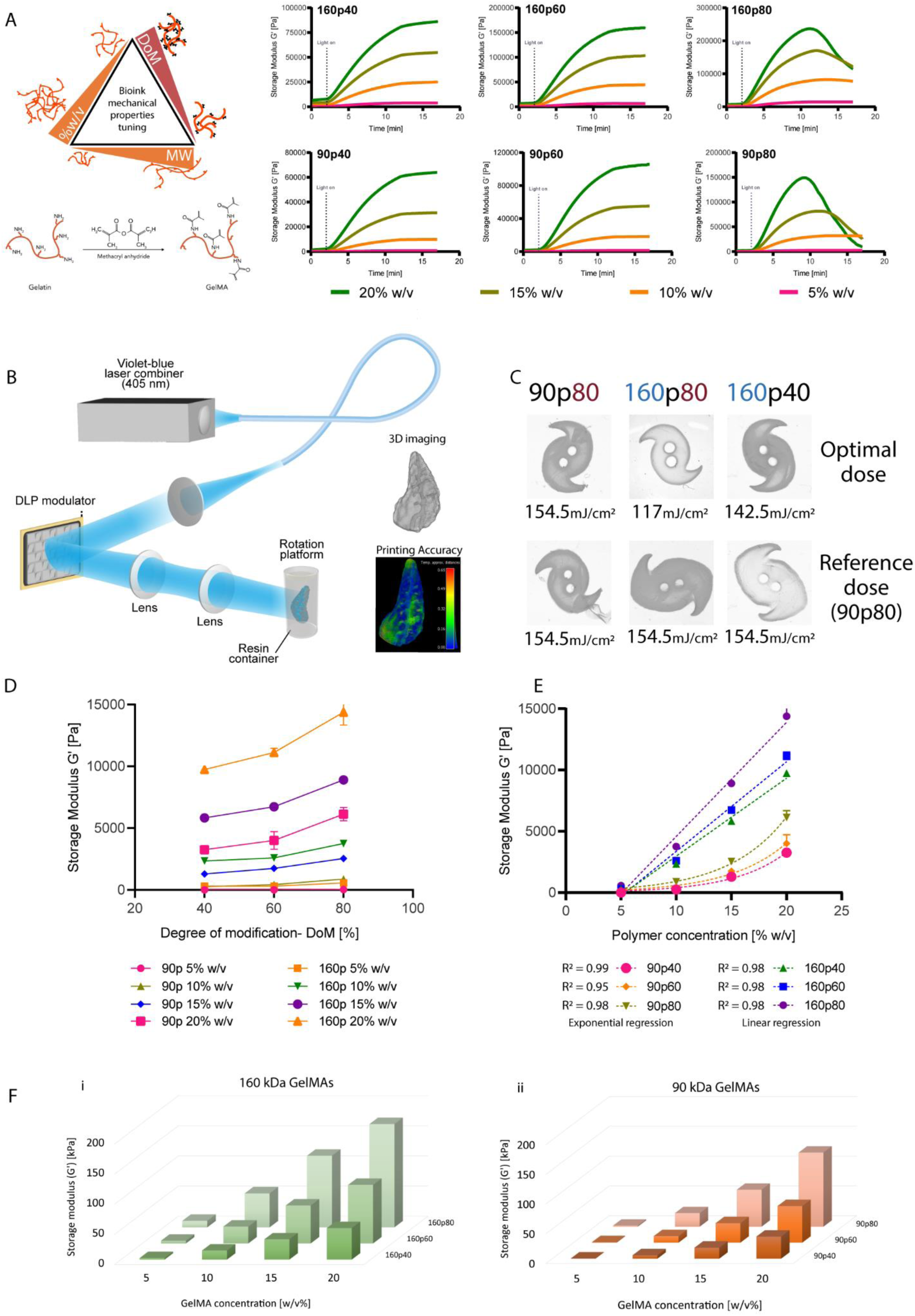
Library of the mechanical properties for the different GelMA hydrogel formulations and how they affect volumetric printing parameters. A) Schematic representation of the GelMA reaction process and overview of the different parameters involved in the design and tuning of the final bioink or bioresin mechanical properties. Representative (n=3) photo-rheology curves for the 90 kDa and the 160 kDa GelMAs with different degrees of modification (40%, 60% and 80%), at different polymer concentrations (5% w/v, 10% w/v, 15% w/v and 20% w/v). B) Experimental setup for computational axial lithography, showing a volumetric printed pancreas model. C) Printing light dose comparison between optimal values specific for the different GelMA formulations, and a reference value (90p80 optimal light dose). D) The hydrogel strengths of the various GelMA formulations at different concentrations. Increased degree of modification leads to greater hydrogel strength. E) The relationship between hydrogel storage modulus and concentration is different depending on the average molecular weight. For the 160 kDa GelMA, with increasing concentration, the storage modulus scales following a linear trend. Conversely, at lower MW (90 kDa), storage modulus scales following a power law, as a function of the concentration (n=3). F) 3D overview of the storage moduli (G’) distribution of the GelMA hydrogels with 160 kDa MW (i) and 90 kDa MW, as determined through photo-rheology (n=3).

Besides reaching high printing accuracy, an optimal printing dose resulted in specific mechanical properties of the final crosslinked constructs. The data shown in **Error! Reference source not found.**D confirms that with increasing DoM the storage modulus of the GelMA hydrogels increases too, in line with previous studies on GelMA-based hydrogels.^[18]^ Similarly, with increasing GelMA concentrations (w/v%) the storage modulus of the hydrogel will increase.

Figure 1E shows how for the 90 kDa GelMAs a power-correlation was established, with R^2^=0.99, 0.95 and 0.98 respectively for GelMA 90p40, 90p60 and 90p80. The 160 kDa GelMAs showed, interestingly, a linear type of correlation R^2^=0.98 for all the 160 kDa GelMAs, with higher molar mass showing increased stiffness. This trend can also be observed at glance, with the 3D representation of the dataset reported in Figure 1F, with the softest gel, 90p40 at 5%, showing a storage modulus as low as 379±47 Pa, and the stiffest gel, 160p80 at 20% showing values of 171.46 kPa. Hence, from these datasets, it can be appreciated how 90p60 and 160p40 GelMAs at 15 % w/v can potentially have similar stiffness values, thus opening opportunities for future mechanistic studies in which cell behavior can be studied in gels with invariant stiffness, but tunable mesh size.

Finally, to maximize consistency and reproducibility of the hydrogels formulations, it is also important to consider the effect photo-initiator and salt concentration. On the one hand, specifically for VBP, higher initiator content also affects light attenuation within the vial, therefore requiring to adjust the incident light intensity to achieve crosslinking, as previously reported.^[3]^ Of particular relevance for processing hydrogels with varying MW, DoM, and polymer concentration, the content of photoinitiator affects the photocrosslinking kinetics, as higher concentrations will lead to faster generation of reactive species upon photoexposure. Moreover, since each hydrogel formulation displays a different amount of reactive groups available for the acrylate kinetic chains to form, maintaining a constant LAP concentration across each experimental group contributes to the differences in gelation kinetics and amount of excess radicals that can potentially damage embedded cells. While from a practical point of view this difference can be accounted adjusting the light dose during printing, strategies that optimize the LAP concentration as a function of the available methacrylate moieties can also be envisioned. A dissertation on how varying the photoinitiator concentration affects the curing of hydrogels with different DoM and MW has been included in the Supplementary Information (**Supplementary Figure S1** and **S2**). Lastly, the ionic content of the aqueous buffer used to produce the bioresin also influences both the gelation kinetics that ultimate gel stiffness, with lower stiffness shown for increasing concentrations of phosphate buffer saline (PBS) in the pre-polymer solution (**Supplementary Figure S3**).

### 2.2 GelMA processed at different temperature leads to different strength, crosslinking rate and printing time

Within the volumetric printing process of GelMA, the thermoreversible behavior of gelatin is particularly advantageous, as bioresin-filled vials are placed at cold temperatures to promote thermal gelation, which prevents cell sedimentation.^[3,19]^ We therefore characterised how the crosslinking temperature could affect the curing kinetics and the optimal printing dose within our volumetric printing set up. Results in Figure 2A showed how processing bioresins at different temperatures (4°C, 21°C, and 37°C) influenced the optimal dose for each selected formulation. Optimal light doses for room temperature photocuring resulted in 165 mJ/cm^2^, 221 mJ/cm^2^ and 255 mJ/cm^2^ for GelMA 160p60, 90p60 and 90p40, respectively. Those values reflected the same trend observed previously, where a denser crosslinking network (e.g., GelMA 160p60), required a lower optimal light dose compared to a less dense network (e.g., GelMA 90p40), even at room temperature. Reference optimal light doses for printing at 4°C (135 mJ/cm^2^, 168 mJ/cm^2^ and 187.5 mJ/cm^2^ for GelMA 160p60, 90p60 and 90p40, respectively), were used as comparison to print each formulation at room temperature, and results showed how a lower crosslinking temperature required a lower optimal light dose.^[19,20]^ Printing at room temperature using optimal doses for the 4°C conditions led to under-polymerized structures, and therefore failure to print.

**FIGURE 2:**
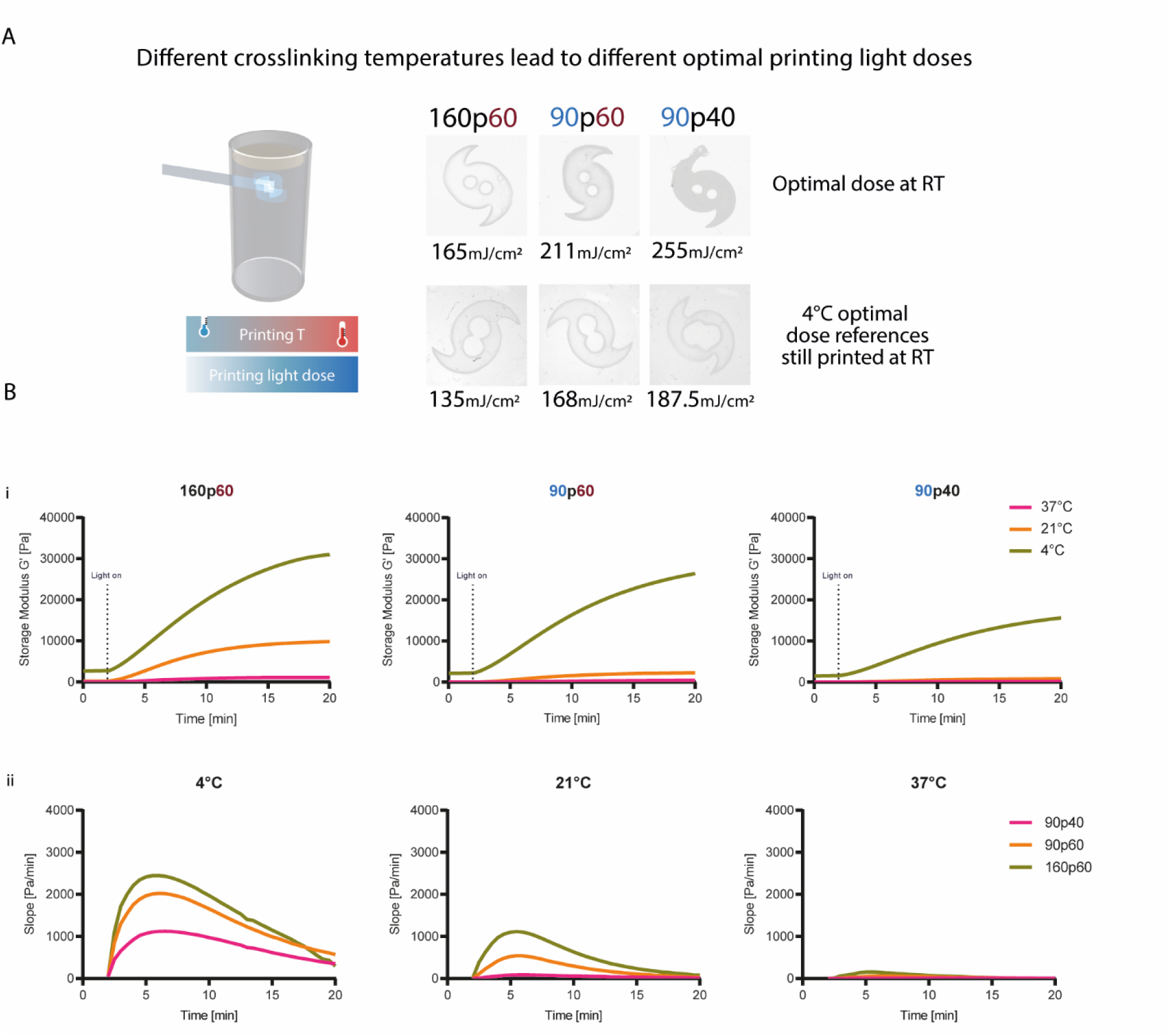
Effect of temperature on volumetric printing parameters, on the final mechanical properties and crosslinking kinetics of different GelMA hydrogels. A) Higher crosslinking temperatures lead to higher optimal printing light doses. Printing light dose comparison between optimal values specific for the different GelMA formulations printed at room temperature, and a reference value (optimal values specific for the different GelMA formulations printed at 4°C). B) (i) Storage moduli (G’) of 160p60, 90p60 and 90p40 at 5 % w/v at 4°C, 21°C and 37° (n=3), showing that photo-curing at decreased temperature results in increased hydrogel stiffness. (ii) Photo-crosslinking rates represented by the first derivatives of the photo-rheology curves of 160p60, 90p60 and 90p40 at 5 % w/v at 4°C, 21°C and 37° (n=3), showing how with increasing temperature, the curing rates slow down. Photorheology was performed with 10 mW cm^−2^, 300 mJ cm^−2^ light source.

To exemplify the thermo-sensitivity of GelMA in hydrogel production, an experiment was designed using 90p40, 90p60 and 160p60, all at a concentration of 5% w/v in 1x PBS. The various GelMA resins were photo-cured and a photo-rheometer was used to showcase the temperature effects in hydrogel crosslinking (Figure 2B). The temperature at which the GelMA resins were cured were 4°C, 21°C (room temperature) and 37°C.

In line with literature, the photo-rheological data presented in Figure 2Bi showed how the hydrogel stiffness, as represented by the storage modulus (G’), is greatly affected by the temperature at which the GelMA is being cured into a biopolymeric network.^[21,22]^ The crosslinking rates, represented by the first derivatives of the photo-rheology curves, are temperature-dependent as well, translating into a faster rates at decreased temperature (Figure 2Bii). In a typical cell printing experiment, the preparation of the bioresin and cell suspension in the prepolymer solution is carried out at room temperature (21°C). Next, regarding volumetric printing, in which the GelMA is cooled to induce physical gelation, the curing kinetics and hydrogel are further affected by the temperature (4°C).^[23]^

### 2.3 Leveraging low MW GelMA to match soft tissues microenvironment mechanical properties

Having elucidated how GelMA based materials result in different final properties depending on the polymer characteristics and on which bioprinting process is adopted, the biological performance of different GelMA from the previously created library was investigated (Figure 3). As proof-of-principle, we aimed for an application in the field of endocrine pancreatic tissue engineering. For this purpose, we bioprinted constructs laden with iβ-cells, an engineered cell line genetically engineered to produce and store insulin in intracellular vesicles (together with the bioluminescent reporter nanoLuc), and to rapidly release it on demand as a response to visible light exposure (475 nm). This enables on demand extracellular delivery of the cargo within the vesicles.^[24,25]^ To select hydrogels within the GelMA library produced with the previous experiments that match the mechanical properties of soft tissues, we focused on GelMA hydrogels at the lowest polymer concentration (5% w/v), and we further investigated their compressive stiffness, and viscoelastic behaviour via a stress-relaxation assay. Figure 3A shows how the shear storage modulus (measured by rheometry) increase for higher molar mass and DoM, and this trend is reflected also in the compressive modulus, as measured in a quasi-static uniaxial compression assay via dynamic mechanical analysis (DMA) (Figure 3B), onsistent with the literature^[21,26]^. More specifically, modulus values of 5% w/v GelMAs (0.1% w/v LAP concentration) ranged from 0.4 ± 0.04 kPa to 0.96 ± 0.1 kPa for the samples 90p40 and 90p80, respectively, while ranged from 1.5 ± 0.18 kPa to 8.9 ± 0.58 kPa for the samples 160p40 and 160p80.

**FIGURE 3:**
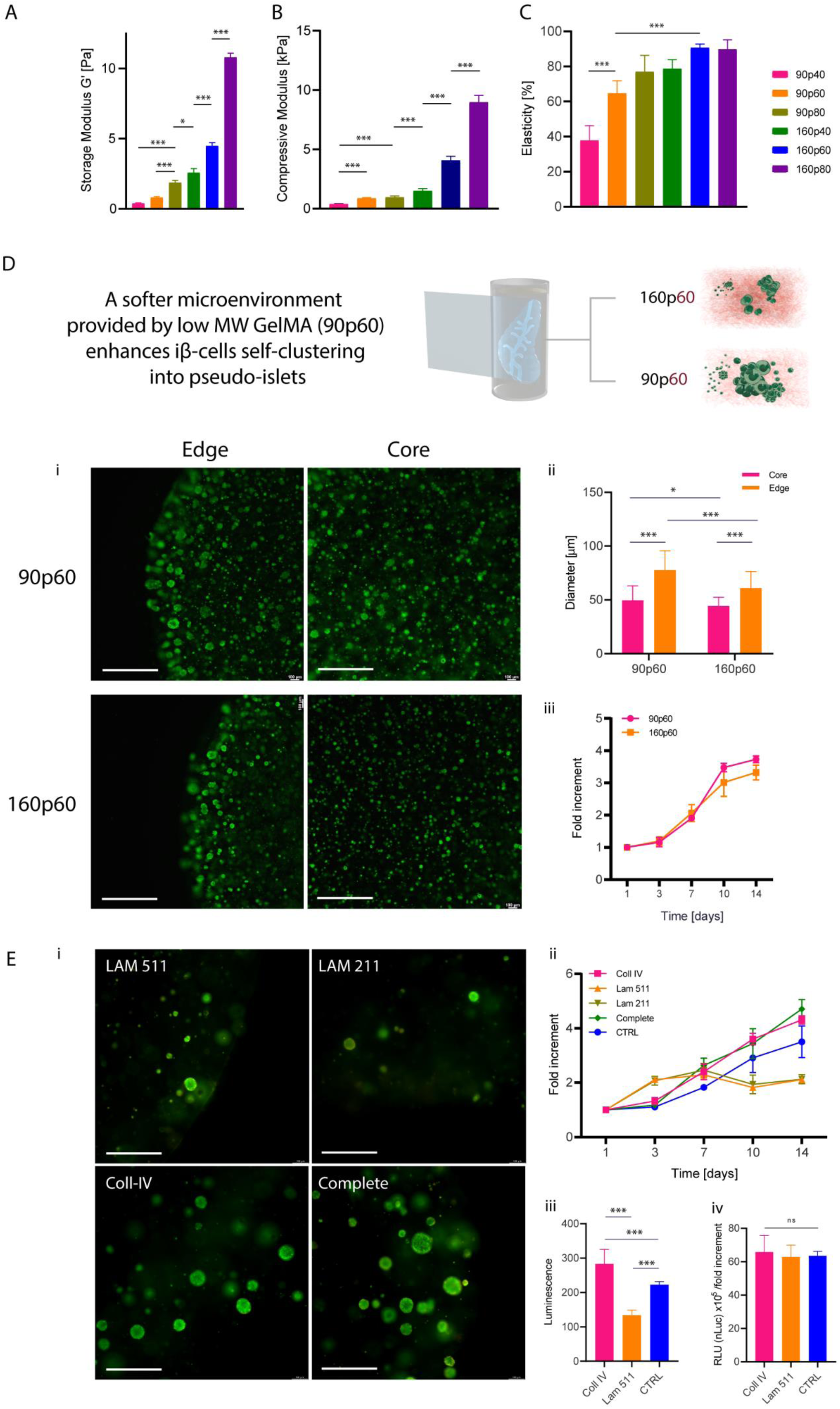
Optimizing 3D cell culture environments via tuning of the GelMA properties via molecular weight and degree of functionalizazion modulation and targeting the mechanical profile of soft tissues. A) storage moduli (G’) B) Compressive mechanical properties and C) elasticity index of the the different GelMA formulations at 5% w/v concentration (n = 4). D) Recapitulating an optimal environment for iβ-cell line leveraging 90p60 GelMA mechanical properties. (i) Effect of molecular weight of GelMA bioresins (90p60 and 160p60) at 5% w/v concentration on (ii) ability to proliferate and form islet-like clusters iβ-cell line and (iii) metabolic activity (n=4). Scale bars: 500µm E) (i) Effect of selected salient macromolecules from the pancreatic ECM on (ii) metabolic activity, (iii) overall and (iv) fold-increment normalised luciferase secretion of iβ-cell line volumetric printed in GelMA 90p60 (n=4). Scale bar: 300 µm

Notably, working with low MW gelatin allowed to modulate the viscoelastic behaviour of the hydrogels. From the stress relaxation curves, an elasticity index (ratio between peak stress, and stress at plateau after relaxation) was extracted (Figure 3C). While high MW gelMA showed a mainly elastic behaviour with slow relaxation (index 89.8% for GelMA 160p80), decreasing MW and DoM allowed to gradually introduce faster relaxation (index 37.8% for 90p40), in a range suitable to mimic several native tissues. To compare the compressive and viscoelastic mechanical properties of the GelMA formulations to soft tissues, we therefore also performed measurements on a porcine pancreatic tissue via mechanical indentation. The compressive properties of porcine pancreas are showed in **Supplementary Figure S4**, over indentation across multiple areas of the tissue. The compressive modulus ranged from 2.3 kPa to 87 kPa, while for the stress relaxation, average the elasticity index ranged from 7 to 34%, after 30 seconds of relaxation, ranges comparable with the lower DoM and MW GelMAs.

An initial biological screening was performed by casting iβ-cells in all the GelMA formulations at 5% w/v and viability was qualitatively tested over 7 days of culture via Live/Dead assay (**Supplementary Figure S5**). A basal membrane extract analogue to Matrigel (commercially available under the name Geltrex) was tested as control. The GelMA 90p40 was unstable in culture, and degraded within the first day of the experiments. For all other samples, from Live/Dead images, cells appeared highly viable both after encapsulation and after 7 days of culture, being homogenously distributed throughout the volume of the hydrogels. The gelMA samples also provided improved stability over degradation compared to the basal membrane extract, which fully degraded by day 7. Considering this results, the 90p60 formulation was selected for further screening on VBP-bioprinted samples, as this formulation displayed low elasticity indexes, and shear modulus and compressive stiffness comparable to the native pancreas^[27–28]^. To reveal the effect of molecular weight on pancreatic cell proliferation and ability to form clusters of pseudo-islets, which is necessary for these cells to functionally secrete insulin,^[9,29]^ 160p60 GelMA was also selected to be compared to 90p60, keeping fixed the degree of modification. Cell metabolic activity and self-clustering capability were therefore tested by comparing volumetric printed gels at 5% w/v, 0.1% w/v LAP embedding 1×10^6^ iβ-cells/mL.

The influence of molecular weight on pseudo-islet cluster formation is shown in Figure 3D. Live/Dead images (Figure 3Di) and quantitative measurements showed how a softer microenvironment enhanced the cell clustering and cluster growth, resulting in significantly larger pseudo-islets in GelMA 90p60, compared to 160p60 (Figure 3Dii). This might be attributed to the lower stiffness and faster relaxation kinetics f GelMA 90p60, due to the shorter polymer chains, which result loose and accessible for cells to self-organise.^[30]^ Moreover, we found pseudo-islets with bigger diameter closer to the edges of the gels in both the formulations, likely due to the shorter diffusion distance for nutrients^[31]^. The overall suitability for cell culture of the hydrogels was quantitatively confirmed by increased metabolic activity values over a period of 14 days, with a 3.7 and 3.3 fold increment for GelMA 90p60 and 160p60, respectively (Figure 3Diii). Having identified the 90p60 formulation as preferred for iβ-cell culture, we further investigated how the addition of ECM components, that are known to be presented in the pancreatic microenvironment, ^[32–36]^ could affect long term cell proliferation and cluster formation (Figure 3E). In particular, collagen type IV, laminin 511, and laminin 211 were chosen because of their well-known contribution to the regulation of islet morphology and survival.^[33,37–39]^ iβ-cells were volumetrically printed in GelMA 90p60 enriched with the single ECM components or with a mix of the three proteins (termed “complete” formulation), and cultured for 14 days. GelMA 90p60 without any additive was used as control. Live/Dead images reported in Figure 3Ei showed overall high viability and self-clustering behaviour in all the formulations, with a qualitatively higher presence of pseudo-islets in the collagen IV and complete formulations, but not in the laminins-only supplemented samples. Further a higher metabolic activity on day 14 for the conditions with collagen IV and the combination of all ECM components was measured (Figure 3Eii), with a fold increment respectively of 4.3 and 4.7 times. Statistically significant differences were found between controls and the collagen IV and complete formulations on day 3, 7, 10 and 14, and between the collagen IV and complete groups vs. the laminins-supplemented conditions at days 10 and 14. These data indicate that the addition of collagen IV is sufficient to enhance iβ-cell proliferation and islet formation, whereas laminin 511 and 211 appeared to reduce proliferation. Finally, to test if the functionalization translates also to improved insulin secretion, we performed a functional assay, with the pristine GelMA controls, and samples enriched with collagen IV and laminin 511. The amount of insulin released by the pseudo-islets was indirectly assessed by measuring the bioluminescent reporter nanoLuc. Results (Figure 3Eiii) showed a significantly higher reporter concentration in the culture medium collected at day 14 in the collagen IV formulation, compared to the control (only GelMA 90p60) and the laminin 511 (resulted as the formulation with the lowest fold increment in metabolic activity). Our findings indicate that this beneficial effect is primarily driven by the improved cell proliferation in presence of collagen IV, rather than in the boosting of the capability of individual cells to produce insulin, as confirmed by the data normalized against the total cell number (Figure 6Eiv). Finally, while our analysis confirms that the 90p60 formulation is suitable to promote the growth and clustering of endocrine cells, we also provided a proof-of-concept indication of the suitability of using low molecular weight GelMAs for supporting the culture of vascular cells, needed for mimicking the vascularized stroma (**Supplementary Figure S6**). Therefore, a co-culture of Human Umbilical Vein Endothelial Cells (HUVECs, GFP-tagged, 2.5×10^6^ cells/mL), with 5×10^6^ hMSCs /mL was photo-encapsulated in a GelMA 90p60 and 160p60 at 5 w/v% concentration. Capillary networks were successfully formed at day 7 only in GelMA 90p60 (**Supplementary Figure S6Ai,ii,iii,iv,v**), compared to the 160p60 formulation (**Supplementary Figure S6Bi,ii,iii**), further confirming how lower molecular weight gelatins provide a more permissive environment for cellular self-assembly across different types of cell lines and applications. It should be noted that no perfusion or interstitial pressure gradients was applied in this time frame. Since these parameters are known to be necessary for promoting capillary vessels maturation and are known to support vascular lumen opening and long term stability in culture,^[40,41]^ future and longer term studies focusing on vasculogenic potential could include dynamic culture settings.

### 2.4 Anisotropic and multi-cellular GeMA-based constructs via multi-material volumetric bioprinting

After successfully demonstrating the ability to photo-crosslink the different GelMA formulations, and showing how GelMA 90p60 could be sculpted via volumetric bioprinting while creating a permissive environment for cell culture, we tested the possibility of forming multi-material constructs containing soft materials for cell embedding and stiffer ones to provide mechanical stability over the culture period. To achieve this, we first set up a strategy to volumetrically print multiple GelMA-based materials in a sequential fashion. In order to create models based on GelMAs with different mechanical characteristics, we selected GelMA 160p60 formulation to be sequentially printed with GelMA 90p60 (Figure 4). The design freedom provided by the volumetric bioprinting process allowed the production of models with movable or articulating parts, like the intertwined rigs printed with GelMA 160p60, or complex self-standing models, like the one resembling the pinched fingers “Italian hand” model emoticon, printed with GelMA 90p60 (Figure 4A). To finally test the printing accuracy of the fabricated structures, we performed a detailed quantitative analysis by calculating the differences between the original STL files and the printed models (**Supplementary Figure S7**). For the intertwined rigs, the average deviation from the original STL was 0.28±0.20 mm, while for the Italian hand 0.20±0.14 mm. Having tested the volumetric printability of both GelMA formulations, multi-material printing was subsequently performed via a sequential approach (Figure 4B).

**FIGURE 4:**
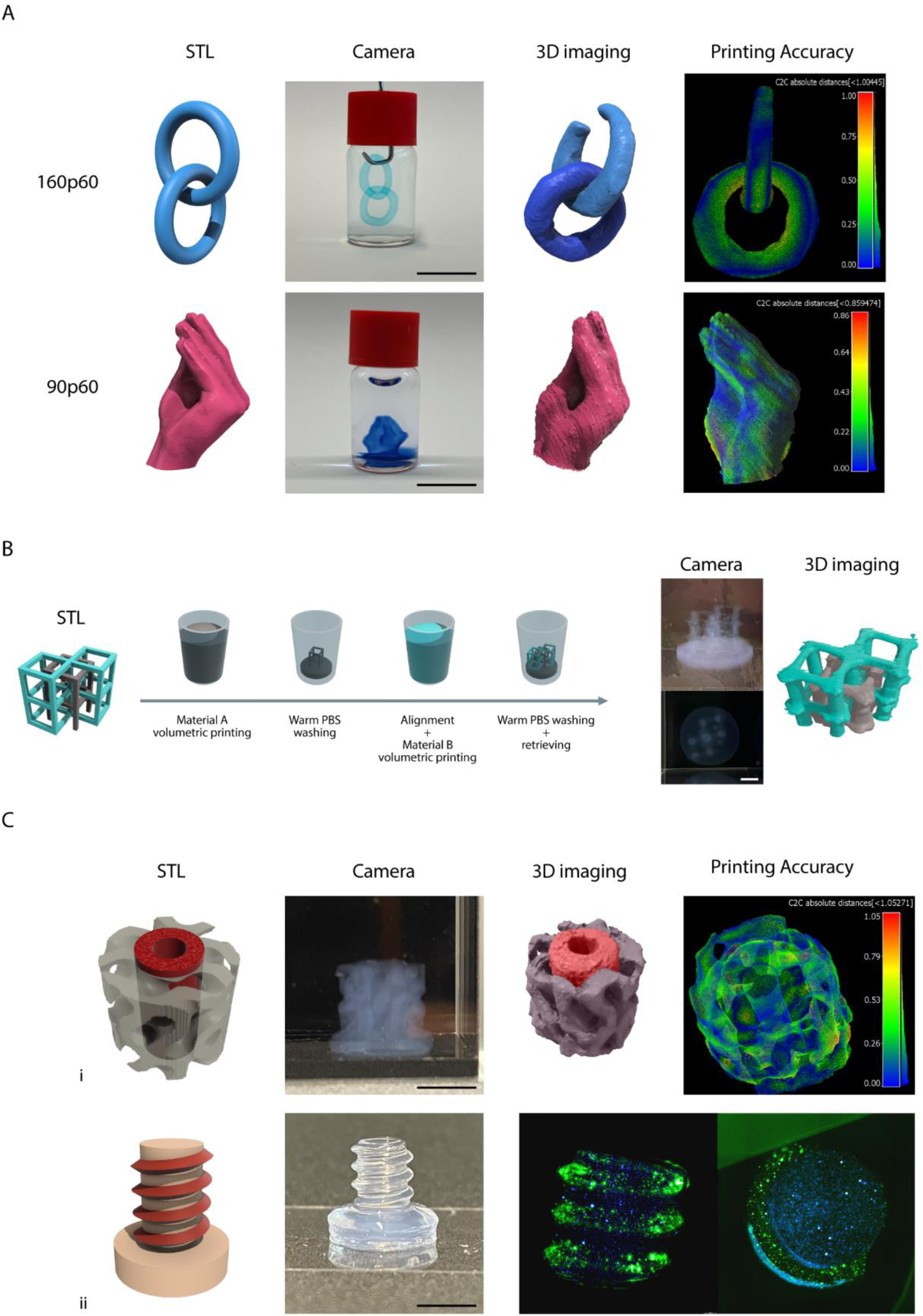
Sequential multi-material volumetric printing. A) Examples of complex structures obtained via volumetric bioprinting of GelMA formulations with different molecular weight, including two interlocked rings, and a model of a hand with pinched fingers. Quantitative printing accuracy 3D maps are provided as a difference between the original STL file and the printed object, showing local variations in the size of the printed features across the whole volume of the objects. Scale bars = 10 mm B) Graphical overview of the sequential multi-material volumetric printing process, and demonstration of two printed independent and intertwined hollow cube scaffolds (digitally coloured in grey and cyan in the 3D light sheet images to facilitate visualization). Scale bar = 3 mm C) Example of multi-material prints: (i) a dual material printed reinforced gyroid (Scale bar = 4 mm) and (ii) a dual material printed screw (Scale bar = 5 mm) with a core and thread printed with formulations embedding iβ-cells with different stainings.

A first volumetric print was performed, then the unpolymerized resin was washed out, and the vial containing the structure printed with the first material was filled with the second material. The alignment of the projections of the model meant to be overprinted with the previously printed structure was performed by leveraging wavelengths (in our case 520 nm) that are orthogonal to the photocrosslinking, which is at 405 nm. This is crucial to prevent unwanted crosslinking, with a subsequent low printing accuracy, if the aligning process was conducted with the same wavelength used to crosslink the bioresin. The second and, eventually, final volumetric printing step was therefore performed.

This approach allowed to print well aligned, and cell-laden objects. For example, in Figure 4Ci, **Error! Reference source not found.** we showed the combination of 160p60 and 90p60 by sequentially printing a reinforced gyroid. The 160p60-based cylindrical scaffold at the core of the 90p60-based gyroid is a possible strategy to exploit the stronger mechanical characteristics of GelMA 160p60 by providing additional stability to the softer GelMA 90p60 structure. Even in this case the printing accuracy analysis showed promising values, with an average deviation from the original STL of 0.25±0.19 mm.

Moreover, to prove the possibility to print multi-cellular constructs on top of anisotropic models, a pancreatic beta-like cell line (iβ-cell line)^[24]^ was stained with different fluorescent membrane dyes, to facilitate visualization, and loaded in distinct GelMA 90p60 solutions.

We showed a sequentially volumetric printed screw, characterised by a 90p60-based core and thread, seeded with the different iβ-cell batches (stained with different fluorescent membrane dyes) (Figure 4Cii). The average deviation from the original screw STL was 0.25±0.24 mm (Supplementary Figure S7D). The herein presented comparison method was not covered before by any other research study for cm^3^-scale complex structures based on soft hydrogels. Supplementary Figure S7 showed a low dimensional error, less a 2%, similar to other studies carried out.^[42,43]^ However, in our study as soft hydrogels were used, it should be kept in mind that these structures tend to swell post printing, therefore introducing a source of dimensional changes prior to the 3D imaging analysis.^[44]^ Overall, these experiments showed that multi-material VBP utilizing materials with different mechanical properties is possible and allows generating both cell-free and cell laden structures. Having proven this potential, this paved the way to next steps investigating how the unique properties of low molecular weight gelatins can be leveraged to enable new high-resolution multi-modal bioprinting techniques converging extrusion and light-based biofabrication.

### 2.5 Low molecular weight gelatins formulations enable facile, plug-and-play embedded extrusion volumetric printing due to their intrinsic self-healing like behaviour

Embedded extrusion bioprinting is a well-established technique in the biofabrication field^[45,46]^ leveraging an ideal support bath for extruding low-viscosity materials.^[46–48]^ In traditional extrusion bioprinting, several artifacts are caused on the overhanging features due to gravity and surface tension-driven effects.^[49]^ To overcome this problem, embedded extrusion printing leverages printing inside of a suspension bath, managing to keep the extruded structure in its predesigned 3D asset and preventing it from collapse. However, the fabrication time increases cubically with the scaling factor, as for all extrusion-based printing techniques, resulting in a disadvantage for cell-laden constructs fabricated at the centimeter-scale.^[3]^

The support bath in which constructs are suspended upon (bio)printing is often complex in formulation, requiring a mixture of multiple materials or lengthy processes and purification steps to obtain minute microgel baths, for example via polymer coacervation,^[50]^ which limits the throughput and broad accessibility of such promising techniques. Alternatively, suspension baths made from bulk polymer gels have been proposes, although these also typically required lengthy preparation steps,^[51]^ or the mixing of multiple components to obtain the proper viscosity^[10]^ and stress relaxation properties. These specifications are needed to allow the extrusion needle to pass through the viscos bath-resin to deposit the ink-material while maintaining the supporting and relax-flow characteristics of the bath-resin.^[52–55]^ Our data, revealed that fluid bulk baths for embedded bioprinting can be produced using only GelMA 90p60, or alternatively low MW unmodified gelatin, without the need of additive, harmful surfactants or rheology modifiers (Figure 5). Our bath-resin, produced using a 5% w/v concentration, upon thermal gelation. Once gelated, 90kDa gelatin showed self-healing like properties which allowed the needle to translate through the bath without creating any scratch or cavity which would affect the bioink deposition quality. The opposite behaviour was observed in 160kDa gelatin, where grooves were created after needle translation (Figure 5A). The same trend was finally observed in the respective GelMAs (90p60 and 160p60). This proved how the modification didn’t affect the possibility to use the low molecular weight formulation as support bath for embedded extrusion printing (**Supplementary Figure S9**).

**FIGURE 5:**
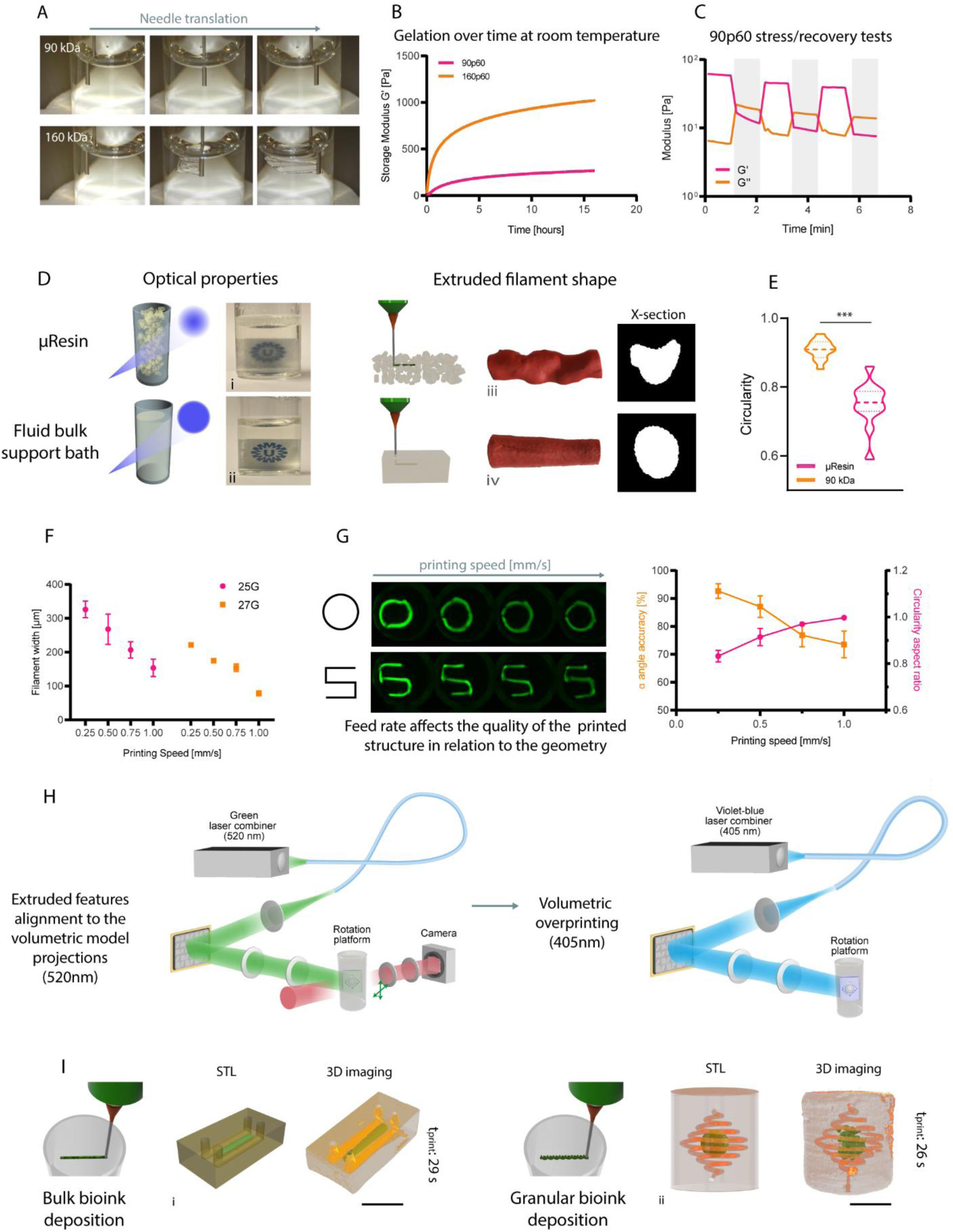
Low molecular weight gelatins enable seamless and omnidirectional extrusion printing as suspension baths, and Embedded extrusion Volumetric Printing (EmVP). A) Fluid bulk support baths mad of 90 kDa gelatin and/or GelMA (see Supplementary Figure S9) show self-healing like properties, whereas 160 kDa GelMA-based bath show slow recovery and mechanical damage caused by the movment of an immersed extrusion nozzle. B) Gelation over time at room temperature of GelMA 90p60 and 160p60. C) 90p60 GelMA rheological properties make it suitable as suspension bath for embedded extrusion bioprinting. Self-healing through low (unshaded, 1% strain, 1 Hz) and high (shaded, 500% strain, 1 Hz) strain cycles (n=3). D) Comparison between μResin and 90p60-based support baths: qualitative characterization of the optical properties (light scattering) of the microparticle-based bioresin (i) and of the fluid bulk support bath (ii). Qualitative characterisation via 3D reconstruction and cross-section visualisation of the embedded filaments extruded in the microparticle-based bioresin (iii) and in the fluid bulk support bath (iv). E) Quantitative characterisation of the circularity of the cross-section from the embedded filaments (n=12). F) Printing accuracy of the two formulations, calculated measuring the filament width as a function of the nozzle diameter and printhead translational velocity (n=5). G) Angle accuracy and circularity aspect ratio quantification at different printing speeds of the embedded extruded filament in 90p60 fluid bulk support bath (n=4). H) Multi-wavelength approach for calibration and printing during EmVP. 520 nm green light, far from LAP excitation spectrum, is used for the manual alignment of the vial in the Z axis and XY plane, in order to match the position of the extruded features with the initial angle at which the vial will start to rotate and send, in synchrony, the projections of the object to be overprinted. Subsequently, the volumetric printing process starts using a violet-blue 405 nm laser line. I) Schematic representation of the EmVP process showing the fabrication of a miocrofluidic chip (channels diameter: 1 mm) (i) via bulk bioink extrusion and centimeter-scale complex structure (channels diameter: 500 µm) (ii) via granular bioink extrusion. Scale bars = 4 mm.

Specifically, 90p60 GelMA is better performing for being used as support bath as the gel stiffness of the physically gelated GelMA is considerably lower (~200 Pa) than 160p60 GelMA (~1000 Pa) after 12 hours of gelation at room temperature (Figure 5B). Most notably, these advantageous properties are stable over multiple hours, facilitating bath storage and performing long sessions of experiments, in contrast with data reported for commonly used gelatins and GelMA, which display flow properties ideal for bioprinting only over a short window of time and temperature during their gelation kinetics.^[56]^

Additional characterisation on the 90p60 GelMA were conducted to further study this behaviour. Stress/recovery tests were performed by alternating low (1%) and high (500%) strain at 1Hz frequency and results showed a higher G”, compared to G’, when the higher strain was applied (Figure S9C). This showed how 90p60 went through a solid-liquid transition when a strain is applied, which could be representative of the needle translating through the gel during the extrusion phase, preventing therefore the gel to crack. After the strain was removed, higher G’ values were registered compared to G”, showing a liquid-solid transition of the gel which led to the possibility to support the extruded filament. In addition, a shear strain sweep test was performed, and results supported the previous data, showing a cross point of G’ and G” at 480% strain, resulting in a solid-liquid transition (**Supplementary Figure S10A**). Moreover, a shear thinning behaviour was observed as the viscosity decreased as the shear strain increased and, under the same conditions, the shear stress increased in a nonlinear fashion (Supplementary Figure S10B). This behaviour was observed both in non-modified gelatin and modified (methacrylated) 90p60, thus opening the possibility to easily build suspension baths for bioprinting that can be either used as sacrificial, temporary support, but also covalently crosslinked post-extrusion printing. In particular, photo-responsive suspension baths can be used for <u>Em</u>bedded extrusion <u>V</u>olumetric <u>P</u>rinting (EmVP), to sculpt the suspension bath into virtually any desired 3D geometry encasing extruded features, as previously demonstrated by our group using a granular bath made of GelMA microgels.

However, when the support bath is meant to be crosslinked via light-triggered reactions, the optical properties play a crucial role as well. Compared to a granular one, a bulk bath resulted more transparent and permissive for light to cross the entire volume without having massive scattering effects, caused instead by the presence of gels microparticles in the granular formulations (Figure 5Di, ii). Moreover, the deposited filament within a granular bath showed a coarser and cracked shape, due to the presence of the microgels and their interstitial porosity that template the extruded filament.^[57]^ Additionally, using granular baths, also entails that, when extruding very low viscosity bioinks or cell suspensions, the bioink can partly filtrate into the inter-particle porosity, compromising printing resolution.^[58]^ Conversely a smooth and tight filament can deposited in the low molecular weight gelatin fluid bulk bath (Figure 5Diii, iv). In fact, measurements of the cross sections of extruded filaments in different baths showed a higher circularity (90%) in strands deposited in fluid bulk support baths, compared to the granular one (74%) (Figure 5E), confirming higher shape fidelity of the filament, and thus to higher printing accuracy of the final extruded geometry. This new approach using low molecular weight single component baths allows therefore to easily obtain high resolution and shape fidelity, without the need of complex multi-material blends or phase-separation systems previously described in the literature to achieve such results.

Following the printability optimization, we managed to extrude features of 80 µm with a GG/PEGDA bioink, previously designed and characterised,^[9]^ which is a smaller dimension than the diameter of the smallest needle (27G) utilized in this study (Figure 5F).

Interestingly, the feed rate of the needle affects the quality of the printed structure in relation to the geometry. In particular, curved and round structures benefit from a higher printing speed, while squared structures and corners benefits from a slower printing speed. This relation was quantified by measuring the angle of squared structures and circularity of round structures in relation to the printing speed (Figure 5G). This crucial aspect must be considered when considering extruding more complex structures comprising both rounded and sharp features, where the printing speed must be consequently adjusted in the g-code in relation to the printed feature.

Finally, we used 90p60 as fluid support bath to generate centimeter-scale, geometrically complex constructs, such as microfluidic chips laden with precisely patterned multiple cell types or materials, by converging volumetric and extrusion bioprinting via EmVP, (Figure 3H), (**Supplementary Figure S11**). Briefly, the designed structure meant to be extruded and the structure to be subsequently volumetrically overprinted are designed as separated models. Embedded extrusion printing was initially performed directly in the volumetric printing vial. Then the vial was placed in a custom developed polychromatic volumetric printer. At this point, the multi-wavelength setup of the volumetric printer is used to first align the extruded features with the projections from the DMD of the model supposed to be overprinted, and finally to perform the crosslinking. More specifically, 520 nm light is used for the alignment, far from the excitation spectrum of LAP to avoid unwanted crosslinking, by sending the projections to the vial, followed by the manual adjustment to match the starting angle of the tomographic reconstruction process. Subsequently, the volumetric printing process was initiated using a 405 nm laser line where a series of light-projections, delivered at a specific dose, recreated the pre/designed geometry onto the existing extruded feature. EmVP performed in fluid bulk support bath further enriches reported techniques by sculpting in tens of seconds different microfluidic geometries and complex models in a high throughput way. Finally, the vial containing the printed constructs was heated to 37 °C to melt the unpolymerized 90p60, and the sample was washed with prewarmed PBS. Figure 6Ii shows biomimetic *in vitro* chip with a bioprinted strand and two parallel channels (channels diameter: 1 mm), potentially simulating blood and lymphatic vessel pair in a tumor chip.^[58]^ Moreover, figure 6Iii shows the possibility to extrude in the fluid support bath not only bulk hydrogels but granular bioinks as well, creating microfluidic structures (channels diameter: 450 µm) with particulate and highly porous cores. All these multi-materials complex structures were obtained through EmVP in less than a minute. Overall, by employing a volumetric bioprinting system comprising multi-color optics, was possible to fabricate highly featured multi-material and multi-cellular geometries, enhancing the accessibility of volumetric printing for advanced tissue models.

## 3. CONCLUSIONS

The unique approach of VBP, which combines the principles of tomographic manufacturing with the use of photoresponsive bioresins, offering ultra-fast speed and precision in the fabrication of complex, cell-laden living constructs, has provided new possibilities for large-scale tissue engineering and regenerative medicine. The development of suitable bioresins, with defined and reproducible properties, is however a critical aspect that requires further exploration. The precise tuning of photocrosslinking kinetics is paramount to ensure accurate volumetric prints and avoid unwanted light dose accumulation, which could lead to over-crosslinking and printing artifacts. Furthermore, the possibility of multi-material printing and the production of complex tissues comprising multiple cell populations are significant hurdles that can be overcome to broaden the applicability of VBP. Further expanding the array of approaches that can volumetrically print complex biological structures as multi- or composite materials (i.e., embedding melt electrowriting-produced opaque polymeric meshes), Embedded extrusion Volumetric Printing (EmVP) was recently developed. EmVP, which combines suspended bath bioprinting with VBP, has demonstrated the potential for positioning multiple cell types within a bioresin vat. However, the resolution of this technique is currently limited by the size of the microgels used. The use of gelatin as a sacrificial, thermoreversible viscosity enhancer offers a potential solution to this problem. However, the temperature-dependent shear-thinning behaviour of commercially available gelatins restricts their use. In this study, the engineering of gelatin macromers was demonstrated as promising strategy to overcome these limitations, thereby creating a library of fit-for-purpose materials for VBP, suspended bath printing, and EmVP. By varying the molecular weight and degree of modification of the gelatins, it is possible to produce bioresins with a wide range of characteristics and printability. The discovery of a subset of gelatins and GelMA that exhibit stable shear-yielding behavior over extended periods represents a significant advancement, offering a new suspension medium for in-bath printing with improved resolution. This selection of materials is compatible for utilization in a multi-color VBP printer to efficiently align different models to produce complex multi-material and multi-cellular geometries. In future developments, heterocellular and anisotropic tissue models that can be volumetrically produces in a high-throughput fashion could have far-reaching implications for biomedical and pharmaceutical research, paving the way for the next generation of *in vitro* tissue models.

## 4. EXPERIMENTAL SECTION

### GelMA formulations

Porcine type A gelatins, used as non-modified and modified formulations, were provided by Rousselot (Belgium). The name coding XpY refers as: X=MW, p=porcine and Y=DoM.

### Photo-rheology for the elucidation of the tunability of GelMA hydrogel strengths

The GelMA types (160p80; 160p60; 160p40 & 90p80; 90p60 and 90p40), were dissolved in 22 mL amber vials at concentrations of 5, 10, 15 and 20 % w/v in a PBS-LAP stock-solution, with LAP being lithium phenyl-2,4,6-trimethylbenzoylphosphinate, at a concentration of 1 mg mL-1 in a 1xPBS solution (0.01 M phosphate buffer, 0.0027 M potassium chloride and 0.137 M sodium chloride, pH 7.4, at 25 °C; Sigma, Belgium). The GelMA resins were placed in an oven (45 °C) for 1.5 hour to improve solubility and they were subsequently vortexed. Next, the GelMA resins were placed in a vacuum chamber and vacuum (50 mbar) was applied for 3 minutes until the foam collapsed. The samples were stored at 4 °C, in the dark.

The photo-crosslinking kinetics of the GelMA resins were studied at 20 °C using an Anton Paar MCR 302e modular compact rheometer (Anton Paar, Belgium) that was equipped with an Omnicure S1500 light source, the wavelength of 365 nm could be selected by using a filter. The intensity was set to 10 mW cm-2 as measured at the quartz base-plate using a DYMAX Accu-Cal 50-LED radiometer. The samples were in situ irradiated from the bottom through the quartz plate. A parallel plate setup (Ø 25 mm, gap 0.300 mm) was used. Frequency and strain sweeps were previously recorded and a shear frequency of 1 Hz and an amplitude (strain) of 1 % were selected as they were within the linear viscoelastic range of the GelMA. The storage and loss moduli (G’/G’’) were monitored over time. The UV light was switched ‘on’ at the 2-minute mark and UV irradiation continued for 10 minutes. After UV irradiation the measurement was continued for an additional 5 minutes. The GelMA samples were stored in an oven at 45 °C during the rheology measurements.

### Dynamic mechanical analysis for the elucidation of the tunability of GelMA hydrogel strengths

The GelMA resins were heated to 45 °C for one hour before crosslinking. The GelMA disks (Ø 6 mm, thickness 2 mm) were photo-crosslinked for 5 minutes, using an in-house build curing chamber with a 365 nm 1 mW cm-2 light source, which had an intensity output of <1 mW cm-2, as measured at the array-plate using a DYMAX Accu-Cal 50-LED radiometer. The crosslinked GelMA disks were submerged in 1xPBS in the incubator (37°C; 5 % CO2) overnight. The mechanical properties of the GelMA hydrogels were tested with the Dynamic Mechanical Analyzer (DMA Q800, TA Instruments), and all tests were performed at room temperature. A quasi-static compression (uniaxial, unconfined) was performed, consisting in a load phase at 0.1 N/min up to 0.3 N. The data was exported to MS Excel and the stress-strain curves were used to calculate the compressive modulus (slope of the load curve in the linear range). To assess the viscoelastic properties, a strain recovery measurement was performed at a constant 20 % strain for 2 minutes and then left for recovery for 1 minute, with a preload force of 0.0010 N. The elasticity index was calculated as being the ratio between the recovered stress and the maximal stress under constant strain (value after 2 mins/highest value * 100).

### Rheology on the GelMA-based support bath for embedded printing

Adjusted Bloom test: Gel casting: 30 mL of each GelMA or gelatin was added to a large petri-dish (Ø 150 mm), obtaining a round surface with a thickness around 1.7 mm of each product. The dish was then left at room temperature to physically gel for 16 hours (overnight). After 16 hours, small round sections were cut and transferred to the stainless-steel base-plate of the rheometer, a plate-plate setup was used with a 25 mm plate spindle. A 1.65 mm gap was used. Paraffin oil and an evaporation cap was applied to prevent sample drying. A thixotropy test was applied to follow up on the storage and loss moduli (G’/G’’). The temperature was set to 21°C, with a tolerance of 0.1 °C for 1 min to allow temperature equilibration. The G’ and G’’ were recorded during 16 hours with a frequency of 1 Hz and shear strain of 1 % (2 second per point). The shear rate ramp test was similarly performed. The shear stress and the viscosity were measured during a logarithmic shear rate ramp from 0.01 to 1000 s-1 (2 second per point). The shear stress ramp test was similarly performed, now using the shear strain during a logarithmic shear stress ramp from 0.01 to 1000 Pa (2 second per point). The stress recovery of the GelMA or gelatin was performed using a thixotropy test: similarly, the temperature to 21°C (tolerance 0.1 °C for 1 min to allow temperature equilibration) and the G’ and G’’ were measured during 2 minutes with a frequency of 1 Hz and shear strain of 1 % (2 second per point). Then a high shear rate was applied (500 s-1) to disrupt the gel for 2 minutes (2 seconds per point). After, the G’ and G’’ were again measured for 10 minutes, with a frequency of 1 Hz and shear strain of 1 % (2 second per point).

### Porcine pancreas mechanical characterization

A frozen porcine pancreas (−18°C) was defrosted at 4°C over 48h. The pancreas was used undissected and uncut and subjected to compression testing using a TA.XT plus texture analyzer (Stable Micro Systems Ltd, UK) at room temperature. A 13 mm diameter plunger was used. A trigger force of 0.005N was applied with an indentation depth of 5mm at speed of 0.5mm/s. after reaching the designated depth the force was recorded for an additional 20 s, thereby providing information about the elasticity of the pancreatic tissue. 49 measurements were performed, covering all the area of the pancreas in an evenly distributed fashion (a grid was used). To determine the compressive modulus, the recorded forces need to be divided over the area at which the force was recorded/applied. The diameter of the plunger is known (i.e., Ø 13 mm) hence strength could be calculated.

### Embedded Extrusion printing

Three-dimensional (bio)printing experiments were performed with a R-GEN 100 (RegenHu, Switzerland), using a pneumatic-driven extrusion printhead. For printing evaluation, 3 mL syringes were loaded with 1 mL of Cy-3.5 stained GG/PEGDA bioink (already characterized in a recent work from our lab),^[9]^ two different needles (25G and 27G) and four different printing speeds (0.25, 0.5, 0.75 and 1 mm/s) were tested to characterize the filaments width. Images (n = 5) of S-shape embedded filaments were taken with a confocal microscope (SPX8, Leica Microsystems, The Netherlands) and analyzed with ImageJ.

### Volumetric printing

A commercial Tomolite v2.0 (Readily3D SA, Switzerland) volumetric bioprinter was used to fabricate the GelMA based constructs. The GelMA formulations (cell-free or embedding cells) were poured at 37°C in Ø10 mm cylindrical glass vials and kept cool at 4 °C to ensure thermal gelation. User-designed CAD files were loaded and processed using the Readily3D Apparite software. The average light intensity across all prints was set at 9.9 mW/cm^−2^. After printing, the thermally gelated bioresin was washed with pre-warmed PBS at 37 °C to remove the uncured GelMA from the printed structure.

### Multimaterial Sequential Volumetric Printing

A first print is performed as described above. The unpolymerized resin is washed out, and the vial containing the structure printed with the first material, is then filled with the second material and the the volumetric printing process is repeated. As long as the optical properties of both materials are compatible with the printing process, a second print (overprinting) can be performed, and upon removal of the second, unreacted resin, a multi-material object is formed. To perform this process correctly, ensuring the correct alignment of the first printed object leveraging a different wavelength prior to performing the second print was crucial. To prevent movement and displacement of the first print during the washing steps, we resorted to print an additional base at the bottom of the vial, to anchor the printed object in place. Then, the alignment of the projections of the model meant to be overprinted with the previously printed structure was performed, followed by the second volumetric printing step. The base can finally be removed at the end of the process, by simply cutting it with a razor blade.

### Embedded Extrusion Volumetric Printing (EmVP)

The extruded features and the model meant to be volumetrically overprint were designed and saved separately as STL files. First, a cylindrical vial was loaded with the 90p60 GelMA at 37°C + 0.1% w/v LAP, everything was then gently mixed directly in the printing vial. The vial was left to thermally gelate at room temperature over night, then placed in a R-GEN 100 extrusion printer (RegenHu, Switzerland) and kept in place through a custom printed holder. 3 mL syringes were loaded with 1 mL of (bio)ink and the G-codes of the extruded features were manually written. Subsequently, the vial was loaded into a custom-made Polychromatic Tomolite volumetric printer (Readily3D SA, Switzerland). To align the extruded features with the model intended to be overprinted, a first series of light-projections was delivered on the rotating vial with a 520 nm laser, orthogonal wavelength to the crosslinking. Therefore, 2D light-projections of the structure meant to be volumetrically printed were delivered with a 405 nm laser, until the curing of the fluid support bath (90p60) was achieved. Finally, the vial was heated to 37°C to dissolve the unpolymerized GelMA and the sample was retrieved and washed with prewarmed PBS.

### Volume comparison for printing accuracy

To visualize and generate the 3D reconstructions of the printed models, GelMA formulations were stained with a fluorescent dye (Cy 3.5 Phosphoramidite), and light-sheet microscopy was used to scan them. Once the 3D-printed structures are manufactured, they are scanned using a customized image system available at our facilities. Different images are obtained once the 3D model is scanned and they are gathered to reconstruct the 3D model using ImageJ. After that, post-processing imaging is carried out to remove outliers and unwanted parts. After that, a detailed quantitative analysis was carried out for calculating the differences between the real STL and the 3D-printed model, using a point-based registration of the meshes. For that, CloudCompare open-source software was used. Both STL files were aligned and the distance between both meshes was computed using the cloud-to-cloud distance, which is the Hausdorff distance algorithm.

### Cells isolation and 2D culture

iβ-cells were obtained and cultured as previously described.^[24]^ Briefly, a 1.1E7 cell line-derived cell clone deficient in glucose-sensitive insulin secretion was transduced with Proinsulin-NanoLuc-derived lentiviral particles and selected in culture medium containing 5 µg/mL blasticidin to create polyclonal INSvesc cells. Next, a monoclonal cell population, was picked based on the best performance for depolarization-triggered nanoLuc secretion. This cell line was co-transfected with the SB100X expression vector pCMV-T7-SB100 (PhCMV-SB100X-pA) and the SB100X-specific transposon pMMZ197 (ITR-PhEF1α-melanopsin-pA-PRPBSA-ypet-P2A-PuroR-pA-ITR) to generate a polyclonal population of iβ-cells that stably expressed melanopsin as well as Proinsulin-NanoLuc cassettes. After selection for two weeks in medium containing 5 µg/mL blasticidin and 1 µg/mL puromycin, the monoclonal iβ-cells were sorted by means of FACS (Becton Dickinson LSRII Fortessa flow cytometer) and screened for blue-light-responsive nanoLuc secretion. iβ-cell were cultured in RPMI 1640 Medium, GlutaMAX™, HEPES (Gibco™, Life technologies) supplemented with FBS (10% v/v) and 1% P/S and used at passage 3-4. cells were cultured in a 95% humidified incubator at 37°C, 5% CO2.

### Statistical Analysis

Results were reported as mean ± standard deviation (S.D.). Statistical analysis was performed using GraphPad Prism 8.0.2 (GraphPad Software, USA). Comparisons between multiple (> 2) experimental groups were assessed via one or two-way ANOVAs, followed by post hoc Bonferroni correction to test differences between groups. Student’s t-test was performed between 2 experimental groups for statistical analysis. Unpaired t-test was used for parametric comparisons of data sets. Non-parametric tests were used when normality could not be assumed. Differences were considered significant when p < 0.05. Significance is expressed on graphs as follows: * p <= 0.05, ** p <= 0.01, *** p <= 0.001, **** p <= 0.0001.

## 5. AKNOWLEDGMENTS

This project received funding from the European Research Council (ERC) under the European Union’s Horizon 2020 research and innovation programme (grant agreement No. 949806, VOLUME-BIO) and from the European’s Union’s Horizon 2020 research and innovation programme under grant agreement No 964497 (ENLIGHT). R.L and J.M acknowledge the funding from the the Gravitation Program “Materials Driven Regeneration”, funded by the Netherlands Organization for Scientific Research (024.003.013). R.L. acknowledges financial support from Dutch Research Council (Vidi, 20387). A.T.O. acknowledges the Basque Government for the postdoctoral fellowship (POS_2021_1_0004).

## SUPPLEMENTARY MATERIAL AND FIGURES

### 1. Influence of the photo-initiator concentration and salt content on photocrosslinking

To correctly set a reproducible hydrogel crosslinking protocol that could be shared, a comparative evaluation of different approaches commonly used in the literature to calculate the LAP photo-initiator concentrations was performed. Three concentrations methods can be discerned from the literature, namely:

1. the stock-solution concentration method (i.e., a standard solution is used for all conditions, e.g., 1 mg/mL of LAP). This is also one of the most used approach in the literature. As a consequence of this method, the ratio between the PI concentration and that of the reactive groups will be different once the GelMA type and concentrations are varied;
2. the mass-ratio concentration method (i.e., the amount of LAP is proportional to the amount of GelMA in the resin, e.g., 1 mg of LAP per 1 g of GelMA);
3. the mol% concentration method (i.e., the amount of LAP is proportional to the methacryloyl content of the GelMA resin, e.g., 2 mol % of LAP relative to the methacryloyl content)

To exemplify what these different concentration methods entail, a numerical example is provided.

#### 1. The stock-solution concentration method

To create a 3 mL GelMA-resin at 5% w/v and 0.1% w/v LAP (1 mg/mL):

| GelMA-type<br>[DoM] | Weight/Volume % | Total volume<br>Resin [mL] | GelMA [mg] | LAP [mg] | Mol% LAP |
| --- | --- | --- | --- | --- | --- |
| 40 | 5 | 3 | 150 mg |  | 48.6 |
| 60 |  |  |  | 0.1 w/v % of | 32.4 |
| 80 |  |  |  | 3 mL = 3 mg<br>LAP | 24.3 |

If a 3 mL GelMA-resin at 10% w/v and 0.1% w/v LAP (1 mg/mL) is produced:

| GelMA-type<br>[DoM] | Weight/Volu<br>me % | Total<br>volume<br>Resin [mL] | GelMA [mg] | LAP [mg] | Mol% LAP |
| --- | --- | --- | --- | --- | --- |
| 40 | 10 | 3 | 300 mg | 0.1 w/v % of | 24.3 |
| 60 |  |  |  | 3 mL = 3 mg | 16.2 |
| 80 |  |  |  | LAP | 12.1 |

The issue with this approach is that the mol% LAP changes both with degree of modification (DoM) and with w/v% of the GelMA-resin (to correct for w/v% you would need to use 0.2 w/v% LAP for the 10 w/v% GelMA). In terms of curing kinetics, the ability to compare resins with one another is lost.

#### 2. The mass-ratio concentration method

To create a 3 mL GelMA-resin at 5% w/v and 10% w/v using a mass-ratio of 1 to 1000 for LAP to GelMA the variability across w/v% disappears.

| GelMA-type<br>[DoM] | Weight/Volu<br>me % | Total<br>volume<br>Resin [mL] | GelMA [mg] | LAP [mg] | Mol% LAP |
| --- | --- | --- | --- | --- | --- |
| 40 | 5 | 3 | 150 mg | 0.15 mg | 2.43 |
| 60 |  |  |  | LAP (1:1000<br>ratio to | 1.62 |
| 80 |  |  |  | GelMA<br>mass) | 1.21 |

If a 3 mL GelMA-resin at 10% w/v and a 1:1000 ratio LAP:GelMA is applied, the following amounts and mol% are required:

| GelMA-type<br>[DoM] | Weight/Volu<br>me % | Total<br>volume<br>Resin [mL] | GelMA [mg] | LAP [mg] | Mol% LAP |
| --- | --- | --- | --- | --- | --- |
| 40 | 10 | 3 | 300 mg | 0.30 mg | 2.43 |
| 60 |  |  |  | LAP (1:1000<br>ratio to | 1.62 |
| 80 |  |  |  | GelMA<br>mass) | 1.21 |

The variability across w/v% of GelMA disappears. However, variability across DoM still exists.

#### 3. The mol% concentration method

To create a 3 mL GelMA-resin at 5% w/v and 10% w/v using the mol% concentration method allows for a comparison across w/v% of GelMA at their respective DoM. The amount of photo-initiator is correlated to the amount of methacryloyl-groups.

| GelMA-type<br>[DoM] | Weight/Volu<br>me % | Total<br>volume<br>Resin [mL] | GelMA [mg] | LAP [mg] | Mol% LAP |
| --- | --- | --- | --- | --- | --- |
| 40 | 5 | 3 | 150 mg | 0.124 | 2 |
| 60 |  |  |  | 0.183 | 2 |
| 80 |  |  |  | 0.247 | 2 |

For a 3 mL GelMA-resin at 10% w/v the following LAP amounts are required:

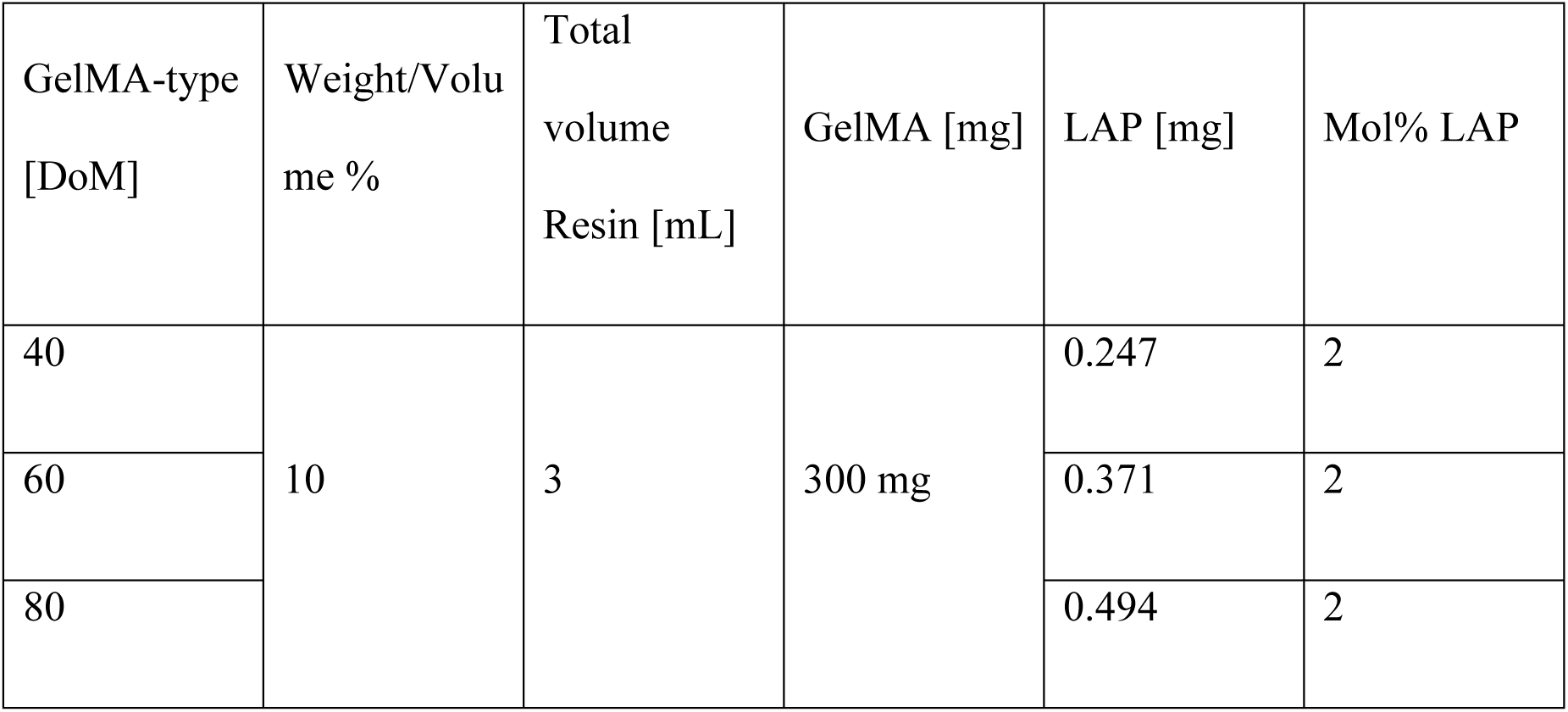

The effect of using the stock-solution concentration method, the mass-ratio and the mol% *photo-initiator* (PI) concentration method in GelMA hydrogel production were studied using photo-rheology (Figure SI-1 and Figure SI-2).

In Supplementary Figure S1 the 90 kDa GelMAs are presented in the left panels, the 160 kDa GelMAs are presented in the right panels. The orange and red curves in Figure SI-1 show the effect of using the PI stock-solution approach. In all graphs photo-crosslinking is completed within the first 2 minutes of UV/vis irradiation (i.e., at an energy dose of 888 mJ). Interestingly, in the 90p40 GelMA condition at a concentration of 5 % w/v, no crosslinking is seen in the mass-to-mass condition (2.43 mol%) or the mol% condition (2 mol%), i.e., an PI amount of 48.6 mol% did incite crosslinking; these data suggest there is an lower limit in which crosslinking can be incited or not (Figure SI-1: panel A).

Furthermore, in all panels of Supplementary Figure S1 it is noted that the light and dark green curves (i.e., indicating the mass-to-mass concentration method) show a delay in curing onset, whereas the light blue and dark blue curves do not show this retarded effect. If this late onset effect was the result of lower PI amounts, than the blue curve in Figure SI-1: panel B should have shown a later curing onset point than the green curve, as the LAP amounts are greater in that variant for the mass-to-mass (2.43 mol%) approach than the mol% approach (2 mol%), however this is not the case. Moreover, the onset in panel C is much later for the 90p60 mass-to-mass approach (1.62 mol%) than for the 90p80 mass-to-mass approach (1.21 mol%), even though the PI amounts are lower in the latter case. Currently it is not clear why this “late curing onset” exists, especially as it is only seen in the mass-to-mass conditions.

**Supplementary Figure S1:**
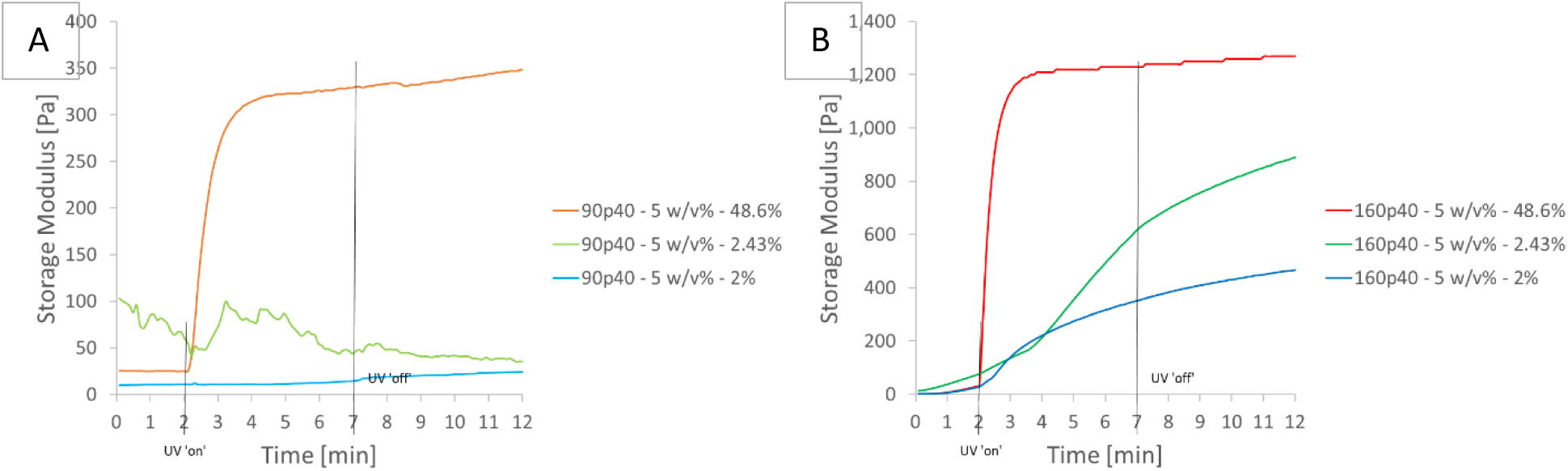
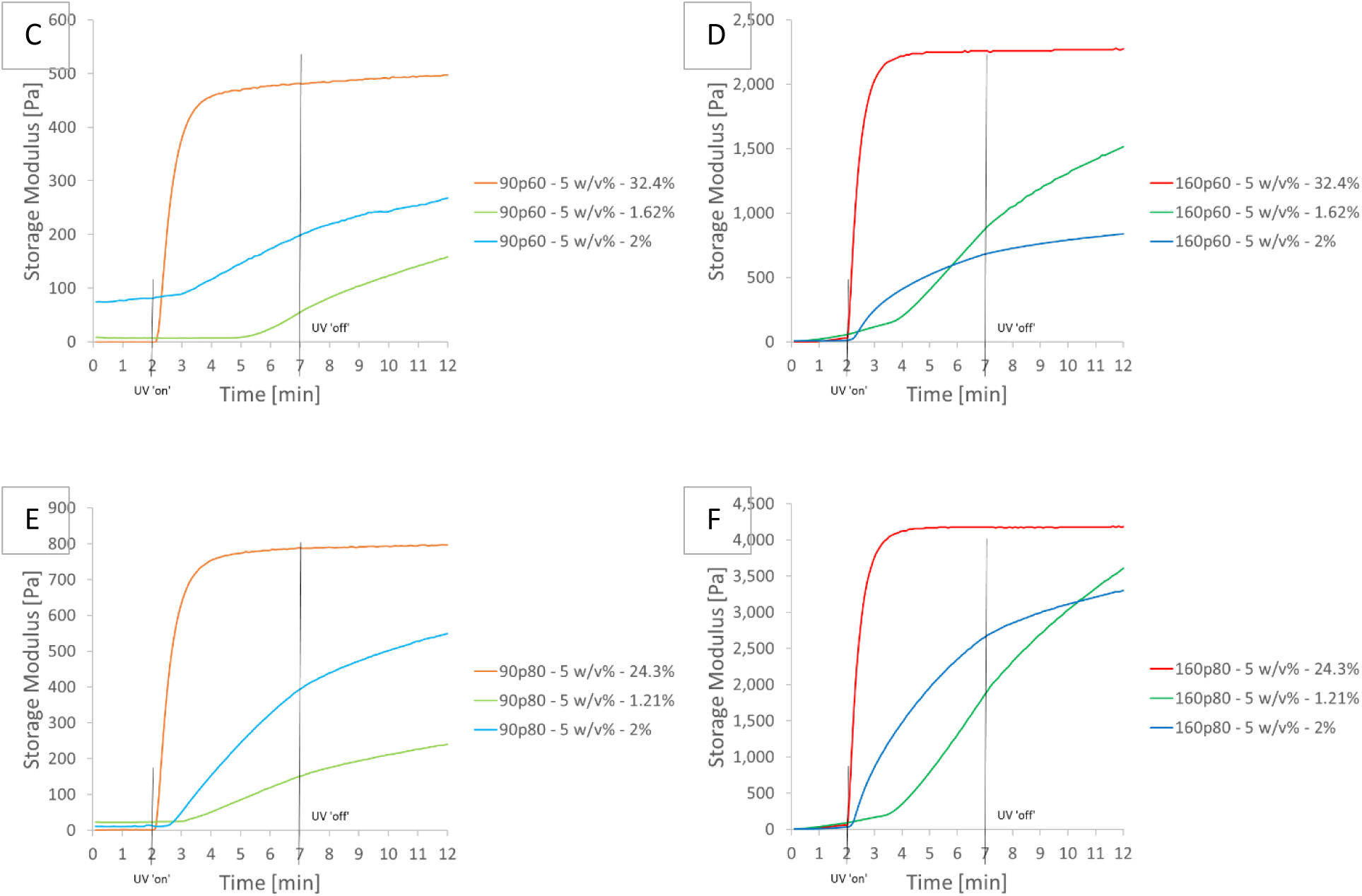
Comparison of storage moduli between using the stock-solution or the mass-to-mass or the mol% concentration methods. The subsequential photo-initiator mol% are given in the graphs. A: GelMA 90p40 at 5 % w/v; B: GelMA 160p40 at 5 % w/v; C: GelMA 90p60 at 5 % w/v; D: GelMA 160p60 at 5 % w/v; E: GelMA 90p80 at 5 % w/v; F: GelMA 160p80 at 5 % w/v.

Supplementary Figure S2 shows the rheology curves of the GelMA conditions at 10 % w/v concentrations. As with Supplementary Figure S1, the stock-solution conditions show fast curing kinetics, with complete crosslinking within the first 2 minutes of UV/vis irradiation. Remarkably, panels B, C, D and F of Supplementary Figure S2, show higher obtained storage moduli for the mass-to-mass PI concentration approach, i.e., the curing kinetics are slower than the stock-solution approach, but the curing is sustained for much longer, resulting in greater hydrogel strength. This effect of slower curing kinetics, but more sustained curing, is most pronounced in the 160p80 condition (Supplementary Figure S2F).

**Supplementary Figure S2:**
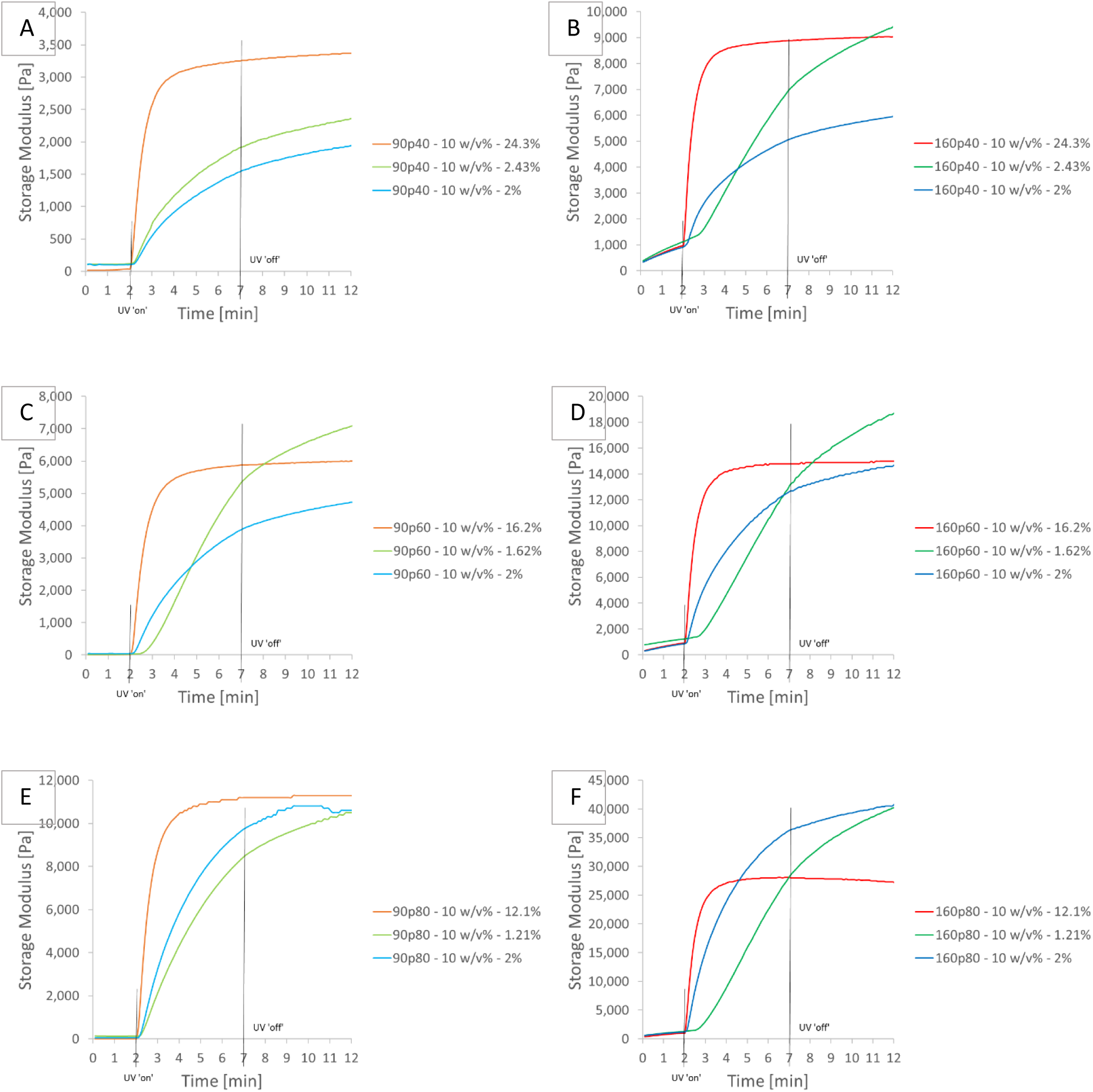
Comparison of storage moduli between using the stock-solution or the mass-to-mass or the mol% concentration methods. The subsequential photo-initiator mol% are given in the graphs. A: GelMA 90p40 at 10 % w/v; B: GelMA 160p40 at 10 % w/v; C: GelMA 90p60 at 10 % w/v; D: GelMA 160p60 at 10 % w/v; E: GelMA 90p80 at 10 % w/v; F: GelMA 160p80 at 10 % w/v.

The importance of describing the applied photo-initiator concentration method in developing GelMA-based bioresins cannot be overestimated with respect to curing kinetics, hydrogel strength, but also debatably, on curing on-set. Three concentration methods were discussed, namely the stock-solution method, the mass-to-mass concentration method and the mol% concentration method. The stock-solution method results in simultaneous variations in w/v% and photo-initiator concentrations, without regard for both the w/v% concentrations of the resins nor the DoM. The mass-to-mass concentration method introduces a fixed ratio between w/v% and photo-initiator (typically 1000:1), but does not take the DoM into account. The mol% concentration method does consider both w/v% and the DoM. The data suggest, there is a minimal photo-initiator amount that is needed to start the crosslinking process, as shown in the 90p40 at 5 % w/v conditions. The data also confirms that with increasing concentration of LAP photo-initiator the curing kinetics will increase, while simultaneously it is shown that a continued increase in LAP photo-initiator concentration leads to a plateau in curing kinetics and even cause a decrease in mechanical strength, i.e., storage modulus of the GelMA hydrogels.

Finally, in Supplementary Figure S3, an example is given using various concentrations of phosphate-buffered saline (PBS) as solution medium for GelMA polymers. PBS is a typical buffer solution used in biological research. It is a water-based salt solution containing disodium hydrogen phosphate, sodium chloride and, in some formulations, potassium chloride and potassium dihydrogen phosphate. Typically, PBS buffers are denoted as 1x or 10x PBS, indicating the concentration levels of the various salts that are present. Supplementary Figure S3, shows that with increasing concentrations of PBS the hydrogel strength decreases, as represented by the storage modulus. By varying the PBS concentration from 0.01x to 1x a decrease of about 60% (from ~13 kPa to ~5 kPa) is shown for the 160p60 GelMA at 5% w/v. The 10x PBS interfered with the photo-curing considerably, the hydrogel produced was not stable, as is clearly visible from the downward trailing of the curve (lightest hue blue curve).

**Supplementary Figure S3:**
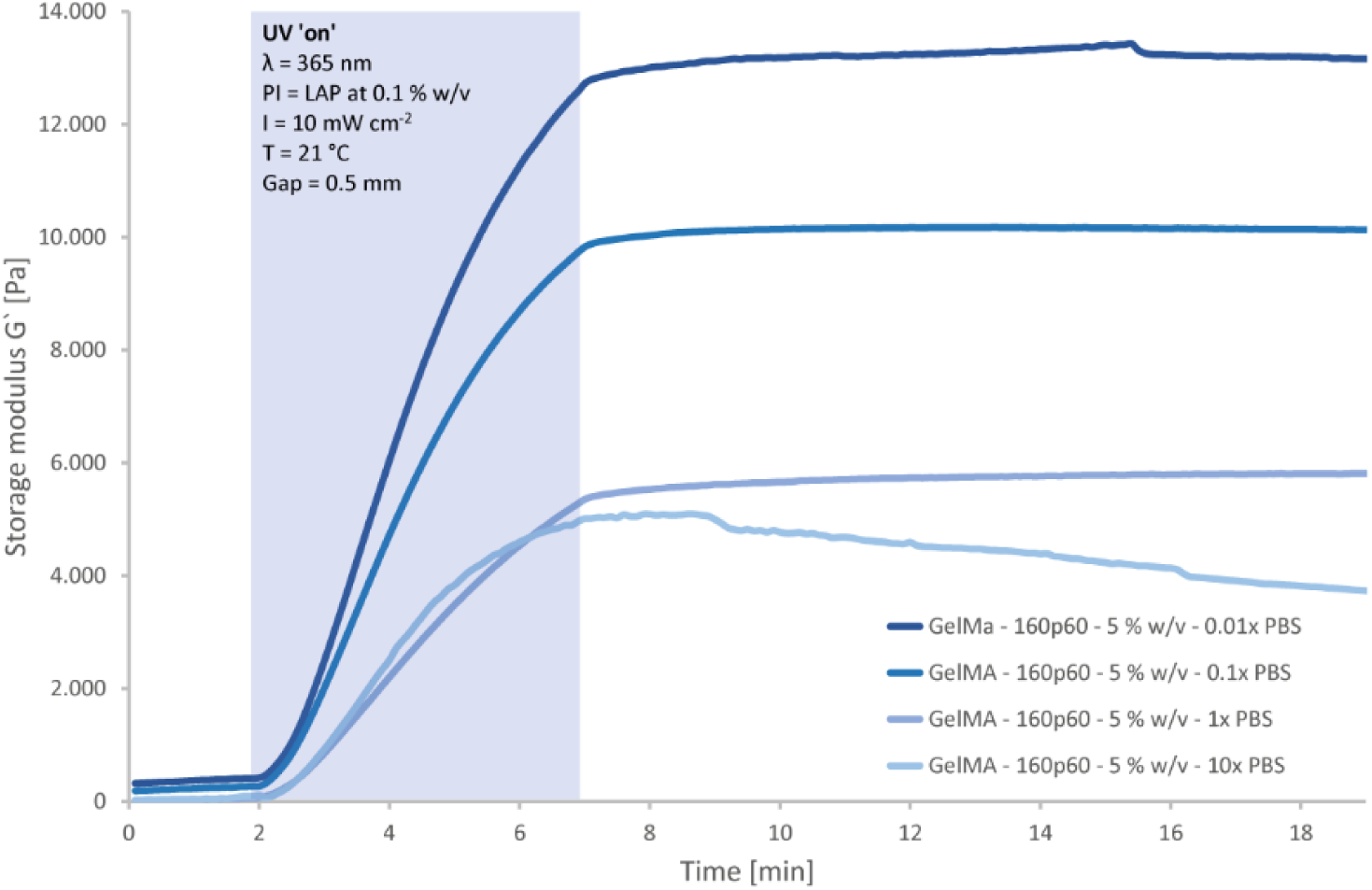
Effect of salt concentration on gelMA mechanical properties. With an increase of concentrations of PBS, the hydrogel strength decreases, as represented by the storage modulus.

#### 2. Cell encapsulation and bioprinting

**Supplementary Figure S4:**
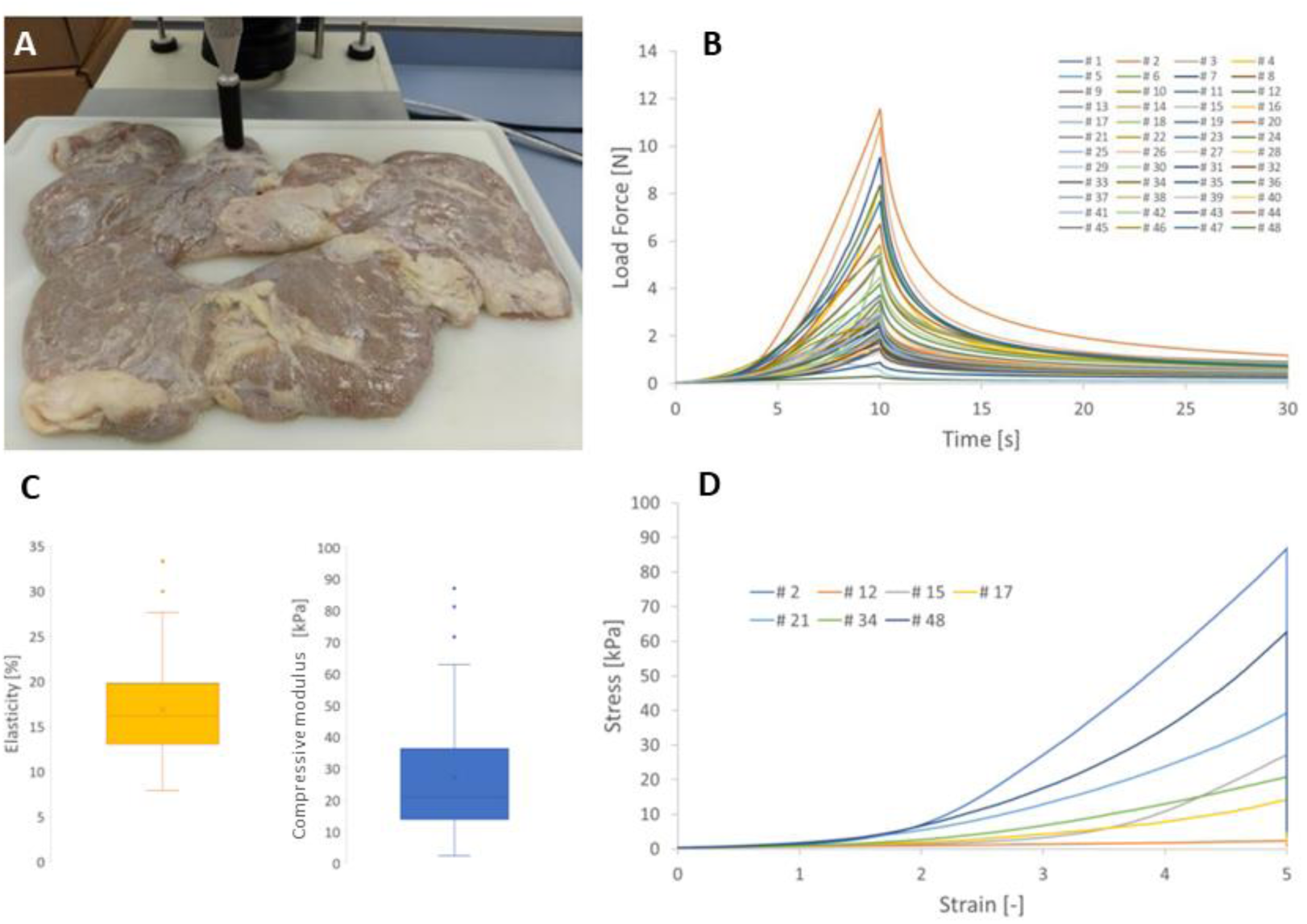
Porcine pancreatic tissue mechanical characterization. A) Porcine pancreas, ready for compression test using an indentation system. B) Recordings of Force as a function of time. The tissue was compressed over a fixed distance of 5 mm after the trigger force of 0.05 N had been exceeded. C) Yellow panel: elasticity of pancreatic tissue. Blue panel: compressive modulus of pancreatic tissue. D) Stress-strain curves of the maximum (#2), minimum (#12), transitions Q1-4 (#17, #21 and #48), average (#15) and median (#34) compression measurements.

**Supplementary Figure S5:**
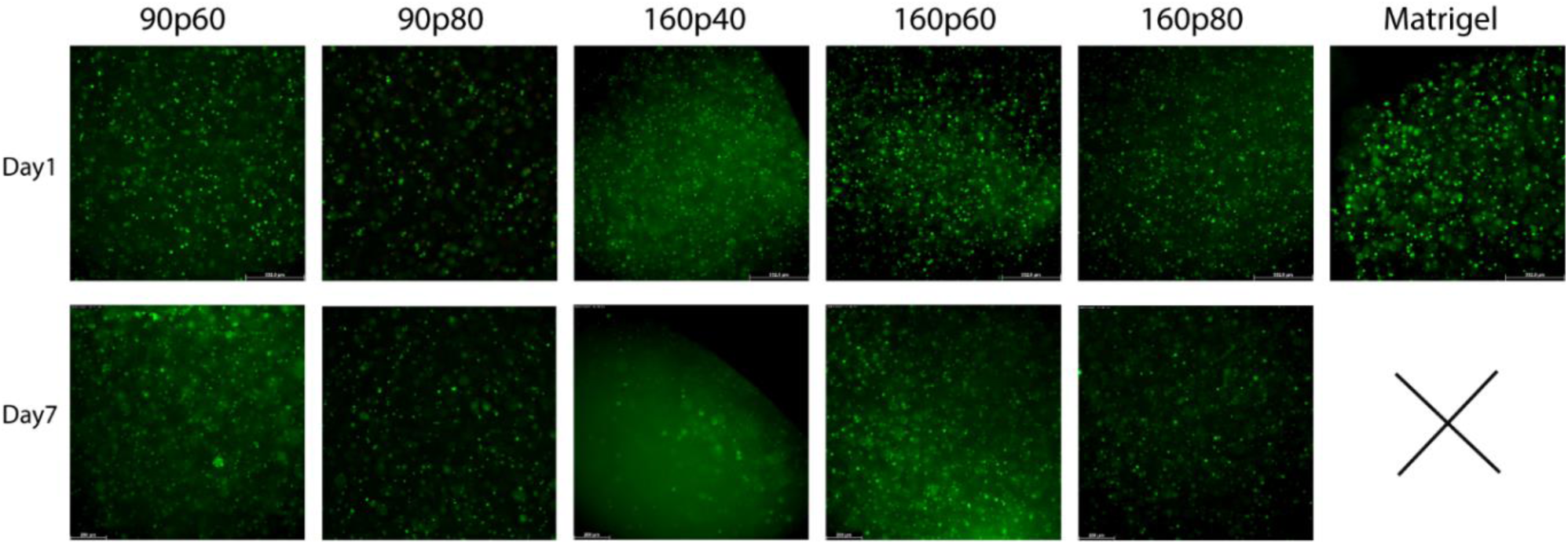
iβ-cells viability screening after embbeding in different gelMA formulations and Matrigel. Live/Dead images showing good cell viability. Geltrex could not be imaged on day 7 as it was degraded. First clusters start to appear after 7 days in GelMA 90p60 Scale bar: 300µm

**Supplementary Figure S6:**
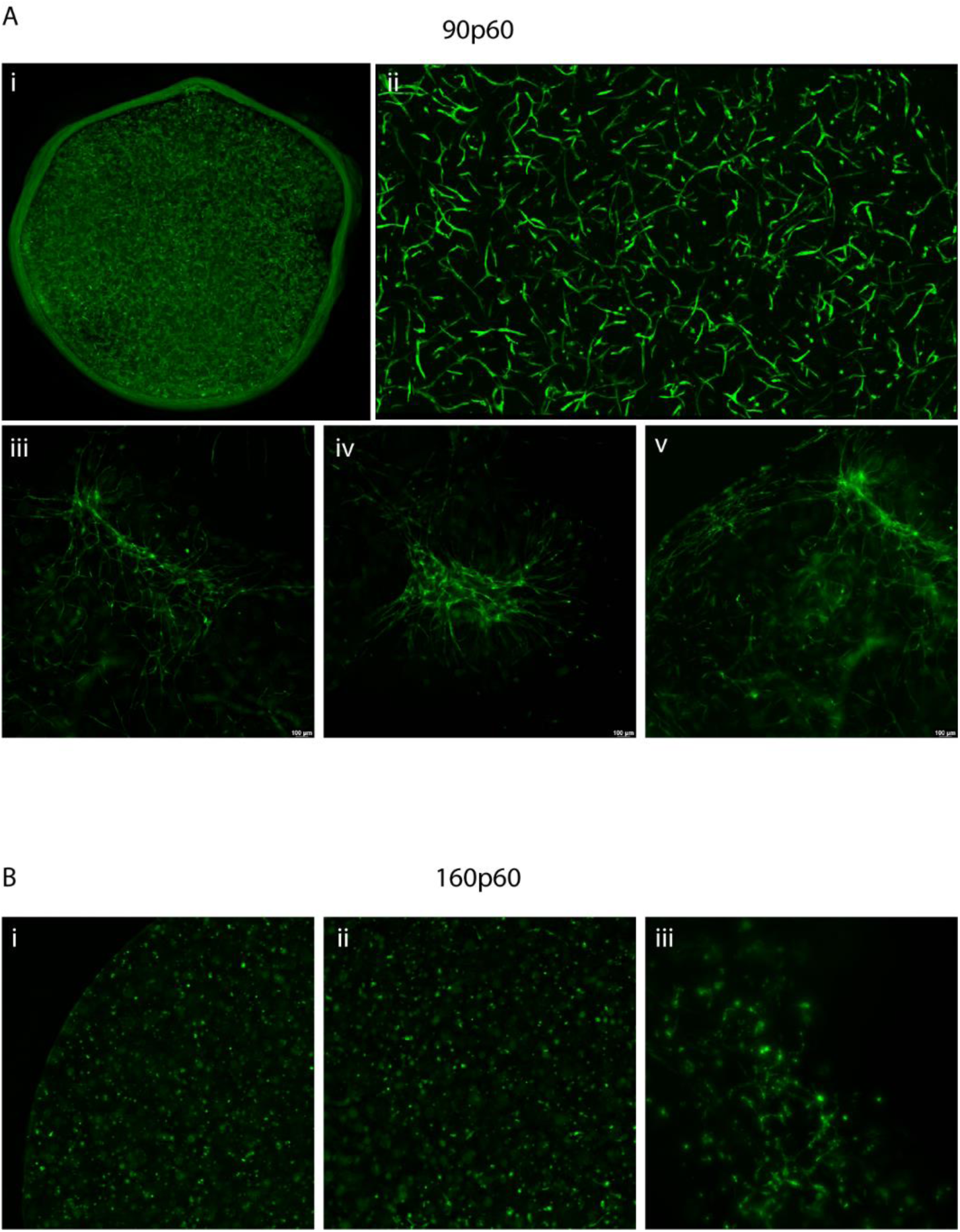
Co-culture of HUVEC and hMSC cells in 90p60 and 160p60 (5%w/v polymer concentration, 0.1% w/v LAP concentration) at day 7. The first capillaries start to appear at day 7. Ai) Disc diameter: 6mm. Zoomed pictures: Aii; Bi,ii : 5x magnification. Aiii,iv,v; Biii : 10x magnification.

#### 3. Printing of complex and multimaterial structures

**Supplementary Figure S7:**
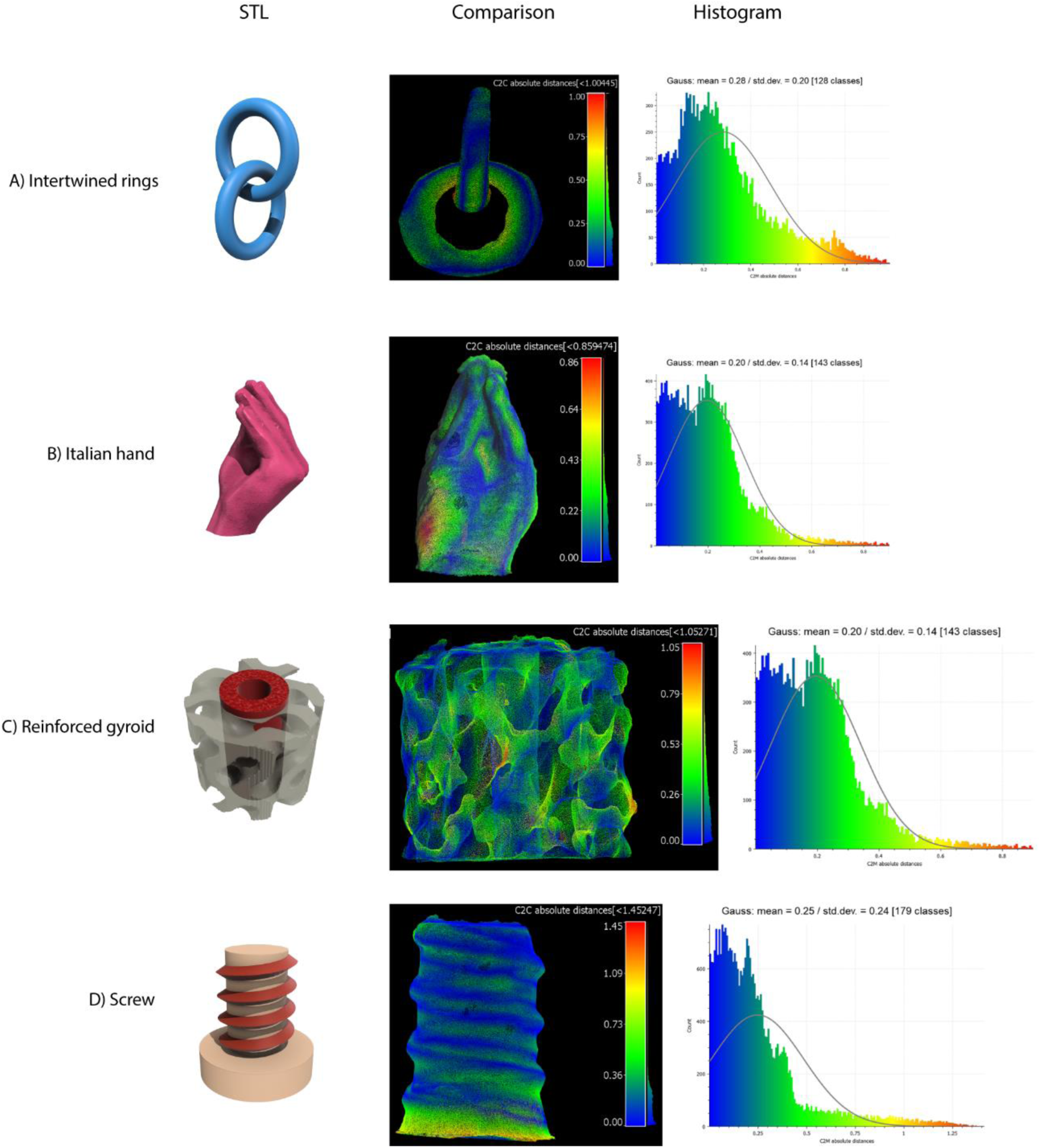
Quantitative printing accuracy. STL models, 3D maps providing the difference between the original STL file and histograms depicting the size variations in mm of the volumetrically printed models: A) intertwined rings, B) Italian hand, C) Reinforced gyroid and D) Screw.

**Supplementary Figure S8:**
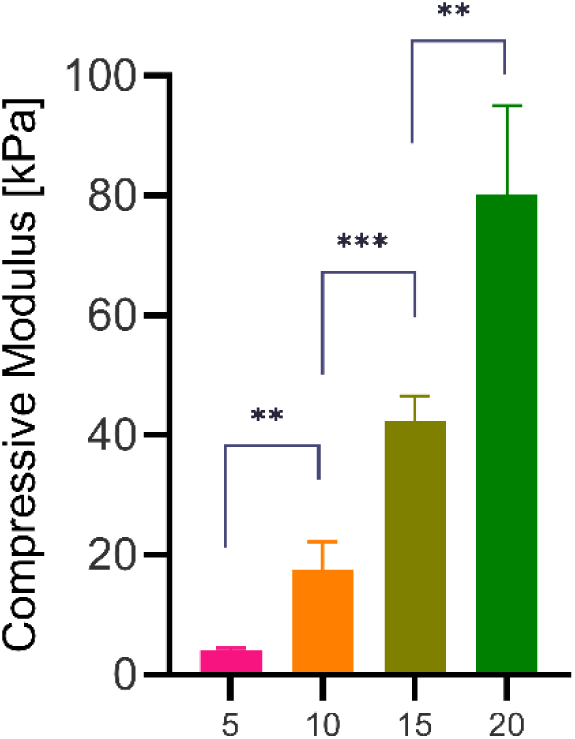
Compressive modulus of gelMA 160p60 at different concentrations. In combination with lower molecular weight gelMA based formulations, 160p60 can act as mechanical support in the multimaterial model.

#### 4. Formation of optimal suspension baths for embedded printing using gelatins at different MW

**Supplementary Figure S9:**
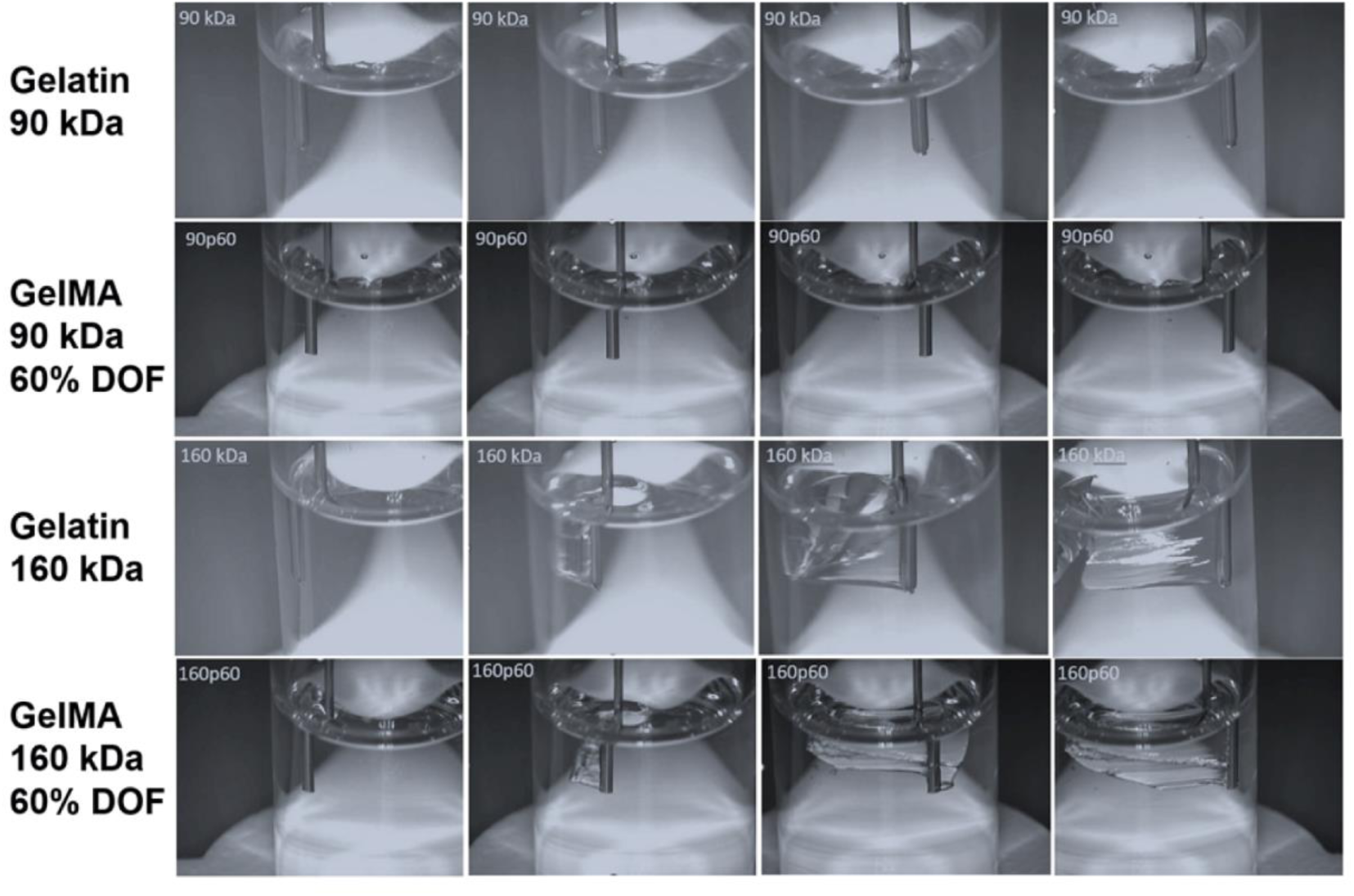
Self-healing like behaviour for both gelatin and gelMA formulations (5% w/v, room temperature). 90kDa gelatin and GelMAs showed self-healing like properties which avoids scratches creation by the needle translations. The opposite behaviour was observed in 160kDa gelatin, where grooves were indeed created. As the same trend was observed both in gelatin and GelMA formulation, it was proved how the modification didn’t affect the possibility to use the low molecular weight formulation as support bath for embedded extrusion printing.

**Supplementary Figure S10:**
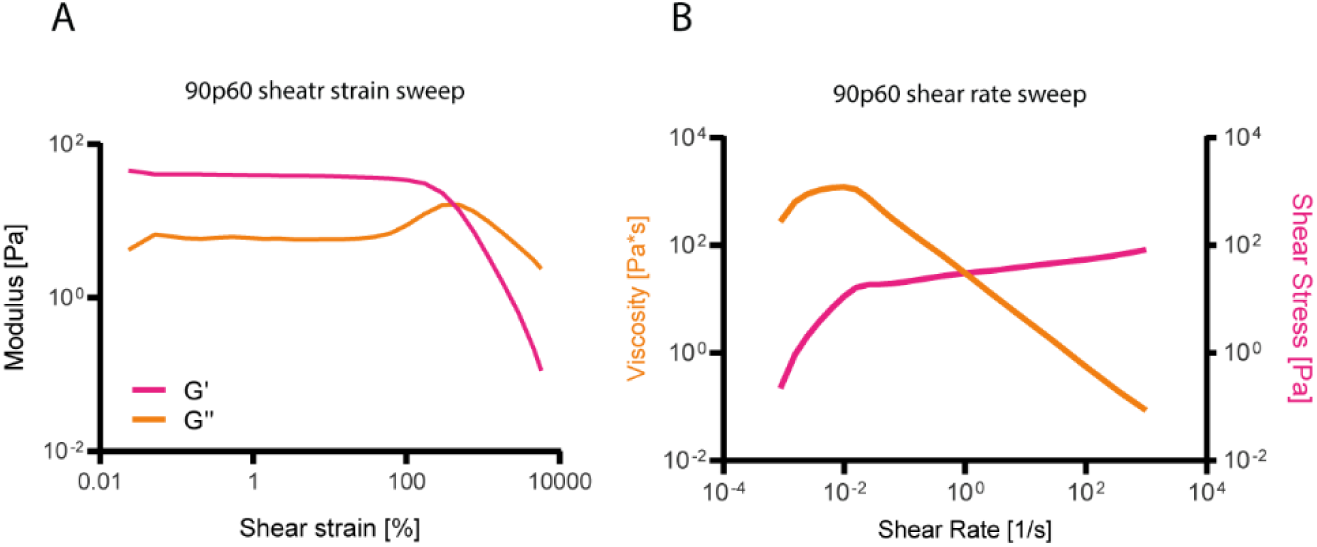
90p60 gelMA rheological properties make it suitable as suspension bath for embedded extrusion bioprinting. A) 90p60 shear-yielding with increase in strain (0.037– 1000%, 1 Hz) (n=3). B) Shear thinning behaviour observed as the viscosity decreased as the shear strain increased and, under the same conditions, the shear stress increased in a nonlinear fashion.

#### 5. Multi-wavelength volumetric bioprinter

**Supplementary Figure S11:**
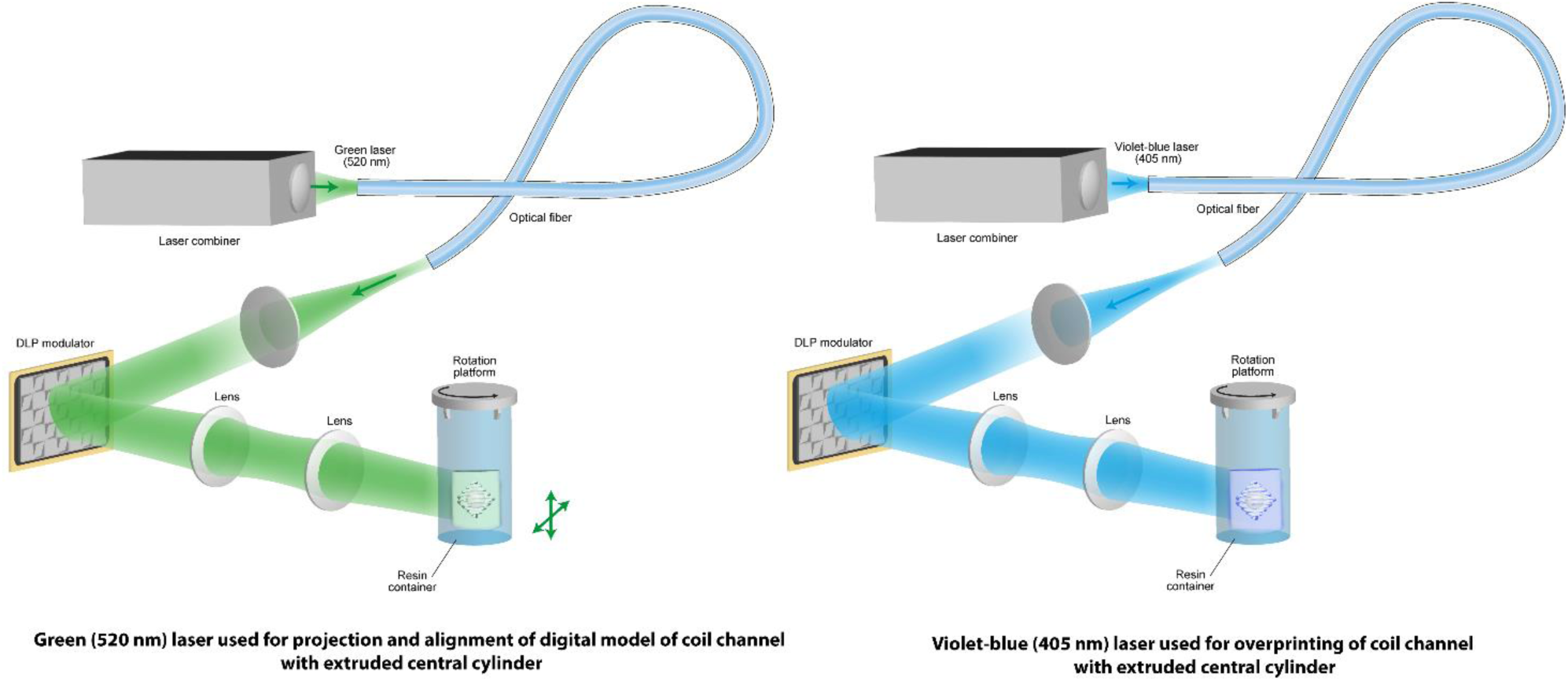
Multi-wavelength approach for calibration and printing during EmVP. Green light, far from LAP excitation spectrum, is used for the manual alignment of the vial in the Z axis and XY plane, in order to match the position of the extruded features with the initial angle at which the vial will start to rotate and send, in synchrony, the projections of the object to be overprinted. Subsequently, the volumetric printing process starts using a violet-blue laser line.

## REFERENCES

[1] B. E. Kelly, I. Bhattacharya, H. Heidari, M. Shusteff, C. M. Spadaccini, H. K. Taylor, Science 2019, 363, 1075.

[2] D. Loterie, P. Delrot, C. Moser, Nat Commun 2020, 11, 852.

[3] P. N. Bernal, P. Delrot, D. Loterie, Y. Li, J. Malda, C. Moser, R. Levato, Advanced Materials 2019, 31, 1904209.

[4] P. N. Bernal, M. Bouwmeester, J. Madrid-Wolff, M. Falandt, S. Florczak, N. G. Rodriguez, Y. Li, G. Größbacher, R. Samsom, M. van Wolferen, L. J. W. van der Laan, P. Delrot, D. Loterie, J. Malda, C. Moser, B. Spee, R. Levato, Advanced Materials 2022, 34, 2110054.

[5] M. Regehly, Y. Garmshausen, M. Reuter, N. F. König, E. Israel, D. P. Kelly, C.-Y. Chou, K. Koch, B. Asfari, S. Hecht, Nature 2020, 588, 620.

[6] L. Hafa, L. Breideband, L. Ramirez Posada, N. Torras, E. Martinez, E. H. K. Stelzer, F. Pampaloni, Advanced Materials 2024, 36, 2306258.

[7] H. Liu, P. Chansoria, P. Delrot, E. Angelidakis, R. Rizzo, D. Rütsche, L. A. Applegate, D. Loterie, M. Zenobi-Wong, Advanced Materials 2022, 34, 2204301.

[8] K. Melde, H. Kremer, M. Shi, S. Seneca, C. Frey, I. Platzman, C. Degel, D. Schmitt, B. Schölkopf, P. Fischer, Sci Adv n.d., 9, eadf6182.

[9] D. Ribezzi, M. Gueye, S. Florczak, F. Dusi, D. de Vos, F. Manente, A. Hierholzer, M. Fussenegger, M. Caiazzo, T. Blunk, J. Malda, R. Levato, Adv Mater 2023, 35, e2301673.

[10] M. B. Riffe, M. D. Davidson, G. Seymour, A. P. Dhand, M. E. Cooke, H. M. Zlotnick, R. R. McLeod, J. A. Burdick, Advanced Materials n.d., n/a, 2309026.

[11] T. G. Molley, T. Hung, K. A. Kilian, Advanced Healthcare Materials 2022, 11, 2201122.

[12] H. Ko, K. Suthiwanich, H. Mary, S. Zanganeh, S.-K. Hu, S. Ahadian, Y. Yang, G. Choi, K. Fetah, Y. Niu, J. J. Mao, A. Khademhosseini, Biofabrication 2019, 11, 025014.

[13] K. S. Lim, J. H. Galarraga, X. Cui, G. C. J. Lindberg, J. A. Burdick, T. B. F. Woodfield, Chem Rev 2020, 120, 10662.

[14] A. Duconseille, D. Andueza, F. Picard, V. Santé-Lhoutellier, T. Astruc, Food Hydrocolloids 2017, 63, 108.

[15] A. T. Stevenson, D. J. Jankus, M. A. Tarshis, A. R. Whittington, Nanoscale 2018, 10, 10094.

[16] L. L. Hench, J. K. West, Chem. Rev. 1990, 90, 33.

[17] S. Panja, B. Dietrich, D. J. Adams, Angewandte Chemie International Edition 2022, 61, e202115021.

[18] J. He, Y. Sun, Q. Gao, C. He, K. Yao, T. Wang, M. Xie, K. Yu, J. Nie, Y. Chen, Y. He, Advanced Healthcare Materials 2023, 12, 2300395.

[19] G. Größbacher, M. Bartolf-Kopp, C. Gergely, P. N. Bernal, S. Florczak, M. de Ruijter, N. G. Rodriguez, J. Groll, J. Malda, T. Jungst, R. Levato, Advanced Materials 2023, 35, 2300756.

[20] M. Falandt, P. N. Bernal, O. Dudaryeva, S. Florczak, G. Größbacher, M. Schweiger, A. Longoni, C. Greant, M. Assunção, O. Nijssen, S. van Vlierberghe, J. Malda, T. Vermonden, R. Levato, Advanced Materials Technologies 2023, 8, 2300026.

[21] Z. Wang, Z. Tian, F. Menard, K. Kim, Biofabrication 2017, 9, 044101.

[22] A. T. Young, O. C. White, M. A. Daniele, Macromolecular Bioscience 2020, 20, 2000183.

[23] P. Chansoria, S. Asif, K. Polkoff, J. Chung, J. A. Piedrahita, R. A. Shirwaiker, ACS Biomater. Sci. Eng. 2021, 7, 5175.

[24] M. Mansouri, S. Xue, M.-D. Hussherr, T. Strittmatter, G. Camenisch, M. Fussenegger, Small 2021, 17, 2101939.

[25] M. Mansouri, M.-D. Hussherr, T. Strittmatter, P. Buchmann, S. Xue, G. Camenisch, M. Fussenegger, Nat Commun 2021, 12, 3388.

[26] C. Ma, J.-B. Choi, Y.-S. Jang, S.-Y. Kim, T.-S. Bae, Y.-K. Kim, J.-M. Park, M.-H. Lee, ACS Omega 2021, 6, 17433.

[27] A. Rubiano, D. Delitto, S. Han, M. Gerber, C. Galitz, J. Trevino, R. M. Thomas, S. J. Hughes, C. S. Simmons, Acta Biomaterialia 2018, 67, 331.

[28] M. Milojević, J. Rožanc, J. Vajda, L. Činč Ćurić, E. Paradiž, A. Stožer, U. Maver, B. Vihar, Biomedicines 2021, 9, 1415.

[29] G. Alessandra, M. Algerta, M. Paola, S. Carsten, L. Cristina, M. Paolo, M. Elisa, T. Gabriella, P. Carla, Cells 2020, 9, 413.

[30] R. Anand, M. S. Amoli, A.-S. Huysecom, P. A. Amorim, H. Agten, L. Geris, V. Bloemen, Biomed. Mater. 2022, 17, 045027.

[31] G. Basara, X. Yue, P. Zorlutuna, Gels 2019, 5, 34.

[32] A. Peloso, L. Urbani, P. Cravedi, R. Katari, P. Maghsoudlou, M. E. A. Fallas, V. Sordi, A. Citro, C. Purroy, G. Niu, J. P. McQuilling, S. Sittadjody, A. C. Farney, S. S. Iskandar, J. P. Zambon, J. Rogers, R. J. Stratta, E. C. Opara, L. Piemonti, C. M. Furdui, S. Soker, P. De Coppi, G. Orlando, Ann Surg 2016, 264, 169.

[33] F. C. Wieland, C. A. van Blitterswijk, A. van Apeldoorn, V. L. S. LaPointe, Journal of Immunology and Regenerative Medicine 2021, 13, 100048.

[34] A. E. Vlahos, S. M. Kinney, B. R. Kingston, S. Keshavjee, S.-Y. Won, A. Martyts, W. C. W. Chan, M. V. Sefton, Biomaterials 2020, 232, 119710.

[35] K. Jiang, D. Chaimov, S. N. Patel, J.-P. Liang, S. C. Wiggins, M. M. Samojlik, A. Rubiano, C. S. Simmons, C. L. Stabler, Biomaterials 2019, 198, 37.

[36] K. Sigmundsson, J. R. M. Ojala, M. K. Öhman, A.-M. Österholm, A. Moreno-Moral, A. Domogatskaya, L. Y. Chong, Y. Sun, X. Chai, J. A. M. Steele, B. George, M. Patarroyo, A.-S. Nilsson, S. Rodin, S. Ghosh, M. M. Stevens, E. Petretto, K. Tryggvason, Matrix Biology 2018, 70, 5.

[37] A. Llacua, B. J. de Haan, S. A. Smink, P. de Vos, Journal of Biomedical Materials Research Part A 2016, 104, 1788.

[38] S. E. Cross, R. H. Vaughan, A. J. Willcox, A. J. McBride, A. A. Abraham, B. Han, J. D. Johnson, E. Maillard, P. A. Bateman, R. D. Ramracheya, P. Rorsman, K. E. Kadler, M. J. Dunne, S. J. Hughes, P. R. V. Johnson, American Journal of Transplantation 2017, 17, 451.

[39] E. Hadavi, J. Leijten, M. Engelse, E. de Koning, P. Jonkheijm, M. Karperien, A. van Apeldoorn, Tissue Engineering Part C: Methods 2019, 25, 71.

[40] K. Haase, R. D. Kamm, Regen Med 2017, 12, 285.

[41] K. Haase, G. S. Offeddu, M. R. Gillrie, R. D. Kamm, Advanced Functional Materials 2020, 30, 2002444.

[42] J. Gehlen, W. Qiu, G. N. Schädli, R. Müller, X.-H. Qin, Acta Biomaterialia 2023, 156, 49.

[43] A. Orth, K. L. Sampson, Y. Zhang, K. Ting, D. A. van Egmond, K. Laqua, T. Lacelle, D. Webber, D. Fatehi, J. Boisvert, C. Paquet, Additive Manufacturing 2022, 56, 102869.

[44] I. Bhattacharya, J. Toombs, H. Taylor, Additive Manufacturing 2021, 47, 102299.

[45] P. S. Gungor-Ozkerim, I. Inci, Y. S. Zhang, A. Khademhosseini, M. R. Dokmeci, Biomater. Sci. 2018, 6, 915.

[46] A. Colly, C. Marquette, J.-M. Frances, E.-J. Courtial, MRS Bulletin 2022, DOI 10.1557/s43577-022-00348-9.

[47] D. J. Shiwarski, A. R. Hudson, J. W. Tashman, A. W. Feinberg, APL Bioengineering 2021, 5, 010904.

[48] K. Zhou, Y. Sun, J. Yang, H. Mao, Z. Gu, J. Mater. Chem. B 2022, 10, 1897.

[49] A. Schwab, R. Levato, M. D’Este, S. Piluso, D. Eglin, J. Malda, Chem. Rev. 2020, 120, 11028.

[50] A. Lee, A.R. Hudson, D.J. Shiwarski, J.W. Tashman, T.J. Hinton, S. Yerneni, J.M. Bliley, P.G. Campbell, A.W. Feinberg. Science 2019, 365(6452):482–487

[51] J. J. Senior, R. J. A. Moakes, M. E. Cooke, S. R. Moxon, A. M. Smith, L. M. Grover, J Vis Exp 2023, DOI 10.3791/64458.

[52] J. Kajtez, M. F. Wesseler, M. Birtele, F. R. Khorasgani, D. Rylander Ottosson, A. Heiskanen, T. Kamperman, J. Leijten, A. Martínez-Serrano, N. B. Larsen, T. E. Angelini, M. Parmar, J. U. Lind, J. Emnéus, Advanced Science 2022, 9, 2201392.

[53] B. J. Albert, C. Wang, C. Williams, J. T. Butcher, Bioprinting 2022, 28, e00242.

[54] V. D. Trikalitis, N. J. J. Kroese, M. Kaya, C. Cofiño-Fabres, S. ten Den, I. S. M. Khalil, S. Misra, B. F. J. M. Koopman, R. Passier, V. Schwach, J. Rouwkema, Biofabrication 2022, 15, 015014.

[55] Q. Li, L. Ma, Z. Gao, J. Yin, P. Liu, H. Yang, L. Shen, H. Zhou, ACS Appl Mater Interfaces 2022, 14, 41695.

[56] W. Liu, M. A. Heinrich, Y. Zhou, A. Akpek, N. Hu, X. Liu, X. Guan, Z. Zhong, X. Jin, A. Khademhosseini, Y. S. Zhang, Advanced Healthcare Materials 2017, 6, 1601451.

[57] M. Becker, M. Gurian, M. Schot, J. Leijten, Advanced Science n.d., n/a, 2204609.

[58] A. Johnson, S. Reimer, R. Childres, G. Cupp, T. C. L. Kohs, O. J. T. McCarty, Y. Kang, Cel. Mol. Bioeng. 2023, 16, 3.

